# Multi-task learning for single-cell multi-modality biology

**DOI:** 10.1101/2022.06.03.494730

**Authors:** Xin Tang, Jiawei Zhang, Yichun He, Xinhe Zhang, Zuwan Lin, Sebastian Partarrieu, Emma Bou Hanna, Zhaolin Ren, Yuhong Yang, Xiao Wang, Na Li, Jie Ding, Jia Liu

**Author notes:** These authors contributed equally: Xin Tang, Jiawei Zhang, Yichun He.

## Abstract

Current biotechnologies can simultaneously measure multi-modality high-dimensional information from the same cell and tissue samples. To analyze the multi-modality data, common tasks such as joint data analysis and cross-modal prediction have been developed. However, current analytical methods are generally designed to process multi-modality data for one specific task without considering the underlying connections between tasks. Here, we present UnitedNet, a multi-task deep neural network that integrates the tasks of joint group identification and cross-modal prediction to analyze multi-modality data. We have found that multi-task learning for joint group identification and cross-modal prediction significantly improves the performance of each task. When applied to various single-cell multi-modality datasets, UnitedNet shows superior performance in each task, achieving better unsupervised and supervised joint group identification and cross-modal prediction performances compared with *state-of-the-art* methods. Furthermore, by considering the spatial information of cells as one modality, UnitedNet substantially improves the accuracy of tissue region identification and enables spatially resolved cross-modal prediction.

## Introduction

Recent advances in single-cell biotechnology make it possible to simultaneously measure gene expression with other high-dimensional modalities in the same cell^1–3^. For example, the patch-seq technique simultaneously measures cell gene expression and intracellular electrical activity^4^, and the ATAC-seq technique jointly measures cell gene expression and DNA accessibility^5^. Such data provide direct and comprehensive views of cellular transcriptional and functional processes simultaneously for the first time. However, many methods have been proposed for single-modality biological data analysis^6^ and cannot be directly applied to multi-modality data. Compared with single-modal analysis, recent studies have identified two common tasks for multi-modal analysis^7^: (i) identification of biologically meaningful groups from different modalities, enabling a deeper biological understanding of cellular identities and functions for different biological systems, and (ii) cross-modal prediction among different modalities, inferring the information of cells that cannot be easily or simultaneously measured. A method that can simultaneously address these two tasks is highly desired to realize the full potential of multi-modality datasets.

Several methods of multi-modal analysis have been developed to address each task separately. For example, for the joint group identification task, multi-modality data integration methods have been developed to fuse different modality measurements into joint representations^8^. The joint representations are then used to further identify cell types and states or tissue regions via unsupervised or supervised classification methods^8–11^. For the cross-modal prediction task, variations of autoencoder-based neural networks have been developed to predict among different modalities^12–17^.

Compared with these methods, an approach capable of addressing these tasks within a unified framework can streamline data analyses and potentially improve the performance of each task^18,19^. Still, combining multi-modal joint group identification and cross-modal prediction into one framework is challenging for the following two reasons. First, each modality measurement has unique statistical characteristics (*e.g*., heterogeneous distributions and noise levels), requiring different statistical assumptions. While there have been several statistical models for different modalities (*e.g*., gene expression measurements^20–22^), a method that can accommodate unknown distributions of simultaneously measured modalities is still lacking. Second, joint group identification and cross-modal prediction typically require different learning workflows and represent separate objectives. Specifically, on the one hand, the workflow of joint group identification needs to accurately extract a joint representation of modalities and then, based on that, identify the similarity/difference between cells, whereas the workflow of the cross-modal prediction needs to find statistical association across modalities. Designing a unifying framework encompassing both workflows is highly nontrivial and crucial to harnessing the full potential of multi-modal datasets. On the other hand, a strategy to coordinate the objectives of these two tasks when training in a unified framework is needed. The objective of joint group identification is to penalize wrong group assignments of cells, whereas that of cross-modal prediction is to minimize the gap between predicted reconstruction and ground truth measurement. Thus, a strategy to integrate the two objectives needs to be designed to avoid performance degradation.

Here, we introduce a deep neural network that performs joint group identification and cross-modal prediction tasks simultaneously for generic single-cell multi-modality data analysis. We build an encoder-decoder-discriminator structure and train it by alternating between the two tasks.

Specifically, this encoder-decoder-discriminator structure does not presume that data distributions are known; instead, it implicitly approximates the statistical characteristics of each modality^23–25^. In addition, we have found that alternating training between joint group identification and cross-modal prediction improves the performance for each task. We have applied this network to various multi-modality datasets (Fig. 1a), including (i) simulated multi-modality data with ground truth labels^26,27^; (ii) simultaneously measured transcriptomics and intracellular electrophysiology^4,28^ (multi-sensing data), (iii) simultaneously profiled transcriptomics and DNA accessibility^5,7^ (multi-omics data), and (iv) spatially resolved transcriptomics and proteomics^29,30^ (spatial-omics data). The results show superior performances in both tasks, achieving better unsupervised and supervised joint group identification and cross-modal prediction compared with other *state-of-the-art* methods.

**Fig. 1.**
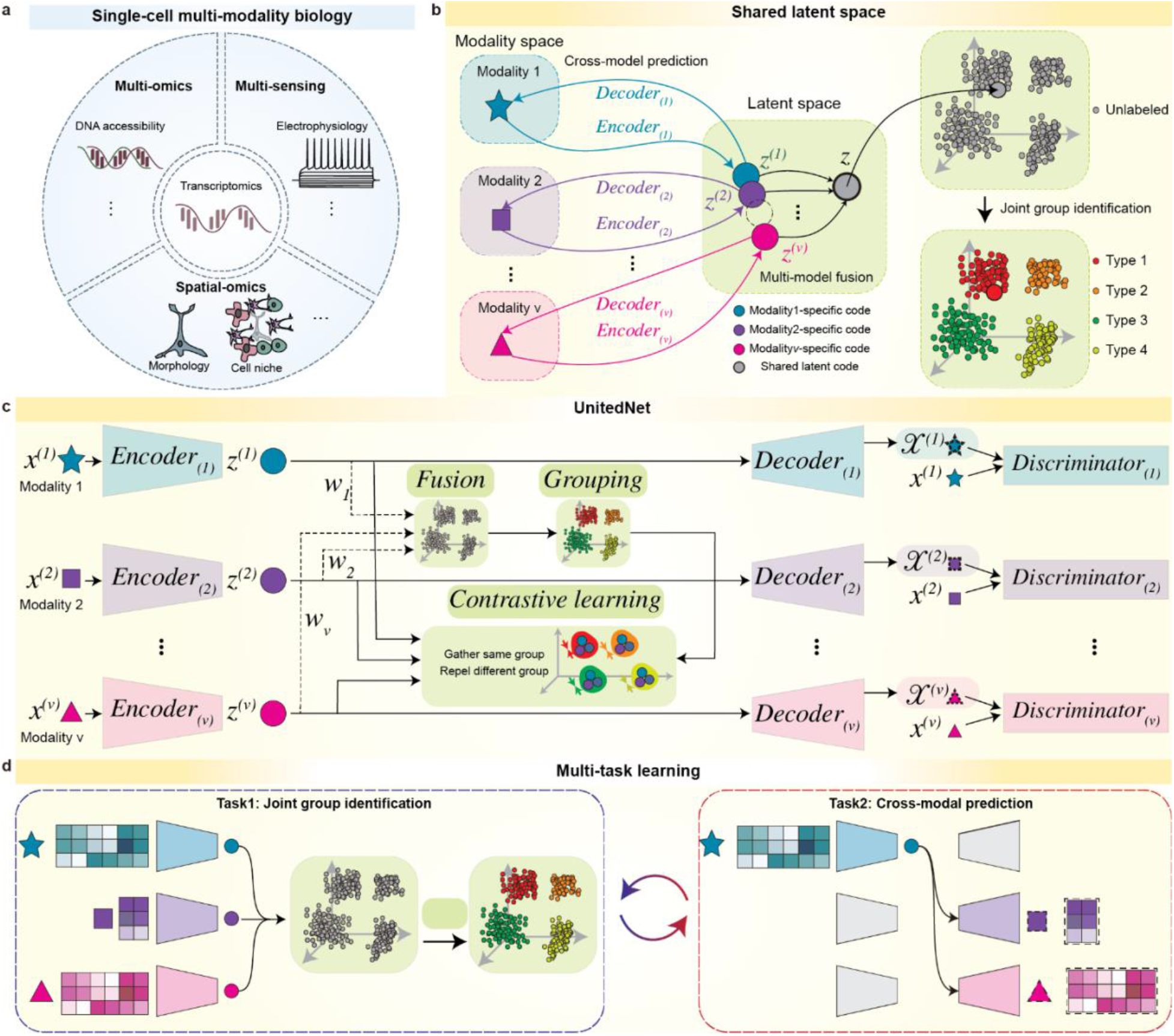
Multi-task learning of single-cell multi-modality biological data by UnitedNet. **a**, Schematics of representative multi-modality biological data: simultaneously measured transcriptomics, intracellular electrophysiology and spatial multi-omics (multi-sensing), (ii) integratively profiled transcriptomics and DNA accessibility (multi-omics) genomics and transcriptomics, and (iii) spatially resolved transcriptomics and proteomics (spatial-omics) data. **b**, Schematics of designation of the shared-latent space. Different modality measurements from the same cell can be projected to a shared latent space as latent codes by encoders. The latent codes can be projected back to the modality space by decoders. In the latent space, latent codes representing different modalities from the same cell can be integrated as a unimodal shared latent code used for joint clustering and group identification. **c**, Schematics showing the network structure of UnitedNet based on the shared-latent space designed in (b). **d**, Schematics showing the multi-task multi-modality data learning by UnitedNet.

## Results

### UnitedNet: a multi-task learning model for single-cell multi-modality biological data analysis

In this paper, we develop UnitedNet, a multi-task learning model to address the challenges discussed in the Introduction. Specifically, for joint group identification, UnitedNet first obtains modality-specific codes (low-dimensional representations) by encoders and then uses an adaptive weighting scheme^31^ to fuse modality-specific codes into shared latent codes (See Methods). Next, it assigns the group labels (*e.g*., cell types) to each cell through unsupervised or supervised group identification networks, where the latter task is also known as annotation/label transfer^8,10,11^ (Fig. 1b and Extended Data Fig. 1a). For cross-modal prediction, UnitedNet first uses the encoder to obtain the source-modality-specific code and then predicts the data of the target modality through the target-modality decoder (Fig. 1b and Extended Data Fig. 1b). The discriminator networks are then trained to distinguish between the true modality and the predicted reconstruction, which competes in an adversarial manner with the training of generative networks, namely the encoders and decoders. Consequently, the discriminator can improve the accuracy of cross-modal prediction^23–25^.

UnitedNet is trained by an overarching loss that consists of (i) an unsupervised clustering loss or a supervised classification loss that separates and closely packs shared latent codes in different clusters to better assign group labels, (ii) a contrastive loss that aligns the different modality-specific latent codes of the same cell and further separates the latent codes from other cells of different clusters, (iii) a reconstruction loss that compares the reconstruction from the encoders and decoders with the original data so that the latent code better represents the cell, (iv) a prediction loss that measures the performance of cross-modal predictions, (v) a discriminator loss that distinguishes the original and reconstructed data in the target modality, and (vi) a generator loss that pushes the decoded data to resemble the original data (Fig. 1c). During the training, we optimized the network parameters by alternately training between the joint group identification and cross-modal prediction tasks and therefore linking their knowledge in the shared latent space (Fig. 1d). A detailed specification of the model, along with a further description of the quantities used in each task, can be found in Methods.

### Validation of UnitedNet performance by ablation analysis

To evaluate the performance of UnitedNet, we used a simulated dataset containing four modalities, DNA, pre-mRNA, mRNA, and protein, with their ground truth labels from *Dyngen*, a multi-omics biological process simulator^26^ (Fig. 2a and see Methods). We first benchmarked the unsupervised joint group identification performance of UnitedNet against standard methods, including (i) *Leiden* for single-modality clustering^32^ and (ii) *Seurat*, which combines weighted nearest neighbor (WNN) analysis^9^ with *Leiden* for multi-modal clustering. UnitedNet exhibits consistently superior unsupervised joint group identification accuracy than the *Leiden* clustering and *Seurat* for different combinations of input modalities (Fig. 2b and Methods). We then performed ablation analysis by removing the cross-modal prediction task in UnitedNet. Notably, we found that without multi-task learning, the unsupervised group identification accuracy significantly decreased (Fig. 2b and Methods). Similarly, we evaluated the cross-modal prediction performance of UnitedNet through ablation analysis. Ablation of either the multi-task learning or the discriminator (termed Dual-autoencoder) substantially reduced the average prediction accuracy of the network. (Fig. 2c, Methods). Together, these benchmark studies and ablation analysis demonstrate the significance of implementing the encoder-decoder-discriminator network structure and multi-task learning scheme for multi-modal data analysis.

**Fig. 2.**
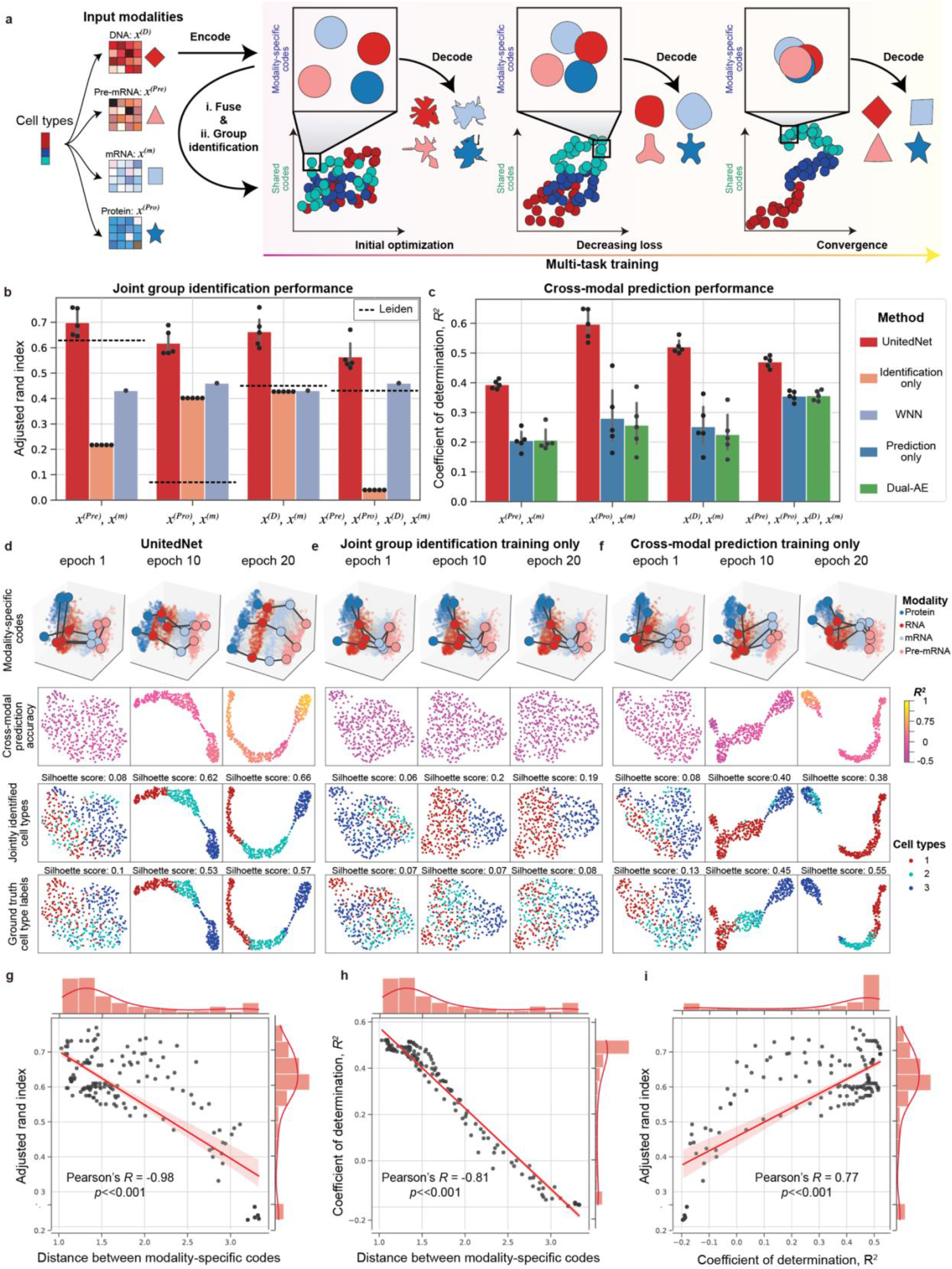
Performance evaluations of UnitedNet on a simulated Dyngen dataset. **a**, Schematics of the optimization procedure of UnitedNet. *x^(D)^, x^(pre)^, x^(m)^*, and *x^(pro)^* represent the simulated modality of DNA pre-mRNA, mRNA, and protein, respectively. Each modality measurement is encoded as a modality-specific code that is then fused as shared-latent codes. The performance of both the joint group identification and the cross-modal prediction task is enhanced as the modality-specific codes are aligned with each other. **b-c**, Barplot reporting the performance comparison of joint group identification (b) and cross-modal prediction (c) of UnitedNet with those from other ablations. The dual autoencoder (dual AE) consists of two vanilla autoencoders without latent space alignment or discriminator. WNN is conducted for the fusion of multiple modalities and then *Leiden* clustering is used for joint group identification on the fused representation. The dashed line in (b) represents the performance of group identification using *Leiden* clustering on the simulated modality of Pre-mRNA, Protein, DNA, and mRNA from *Dyngen*. **d-f**, The latent space visualization by uniform manifold approximation and projection (UMAP)^40^ of UnitedNet (d) and other ablation versions in which we remove the training of cross-modal prediction task (e) or joint group identification task (f) with different training epochs for the *Dyngen* dataset with four modalities. The latent codes are colored to represent different prediction R^2^ values (second row), cell type labels (third row), and ground truth labels (last row). Three representative codes in each of the modalities are highlighted in the first row to show the alignment between modalities. As the training epoch increases, the modality-specific codes are aligned with each other in UnitedNet (d) while in other ablations (e and f) the codes are misaligned. **g**, The correlation plot between the inter-modality distance of modality-specific codes and joint group identification performance. **h**, The correlation plot between the inter-modality distance of modality-specific codes and cross-modal prediction performance. **i**, The correlation plot between cell typing performance and prediction performance.

We then studied why multi-task learning can improve the performance of both tasks. Based on the previous designation of shared-latent space in multi-modal and multi-task learning (Fig. 1b), we hypothesized that the joint training of joint group identification and cross-modal prediction tasks will reinforce each other through the shared latent space (Fig. 2a). Therefore, we compared the shared latent codes learned from single-task training with multi-task training by UnitedNet using the simulated four-modality *Dyngen* dataset. The results show that compared with single-task learning (Fig. 2e, f), multi-task learning better aligned the modality-specific codes and better separated the clusters of the shared codes in the latent space (Fig. 2d). Together, they improved group identification efficiency and cross-modal prediction accuracy upon the model trained with single-task learning (Extended Data Fig. 2). We further quantified the relationship between joint group identification and cross-modal prediction tasks throughout the training procedure. The results show that the performances of both tasks improved as the distance between the modality-specific codes decreased (Fig. 2g, h). Overall, the performances of group identification and cross-modal prediction tasks exhibit a strong positive correlation (Fig. 2i).

In addition, we showed the robustness of UnitedNet when handling datasets with modality-specific noise. In applications, noises arising from different sources, such as the sequencing dropout effect and feature measurement error in multi-modality biological datasets, typically affect the performance of the network. Therefore, we tested the performance of UnitedNet in simulated biological datasets with different controlled noise levels^27^ and simulated multi-modality imaging datasets^31,33^. The results show that the capability of UnitedNet to identify and suppress noise enables it to outperform the *state-of-the-art* models (Extended Fig. 3).

### UnitedNet-enabled multi-modal analysis for multi-sensing data

To demonstrate the ability of UnitedNet to analyze realistic multi-modality biological data, we first applied UnitedNet on a relatively small-scale Patch-seq dataset, which simultaneously characterizes the cell morphological (M), electrophysiological (E), and transcriptomic (T) features from thousands of GABAergic interneurons in the mouse visual cortex^28^. UnitedNet allowed for simultaneous unsupervised joint group identification and cross-modal prediction to identify cell types and predict modality-specific features, respectively (Fig. 3a).

**Fig. 3.**
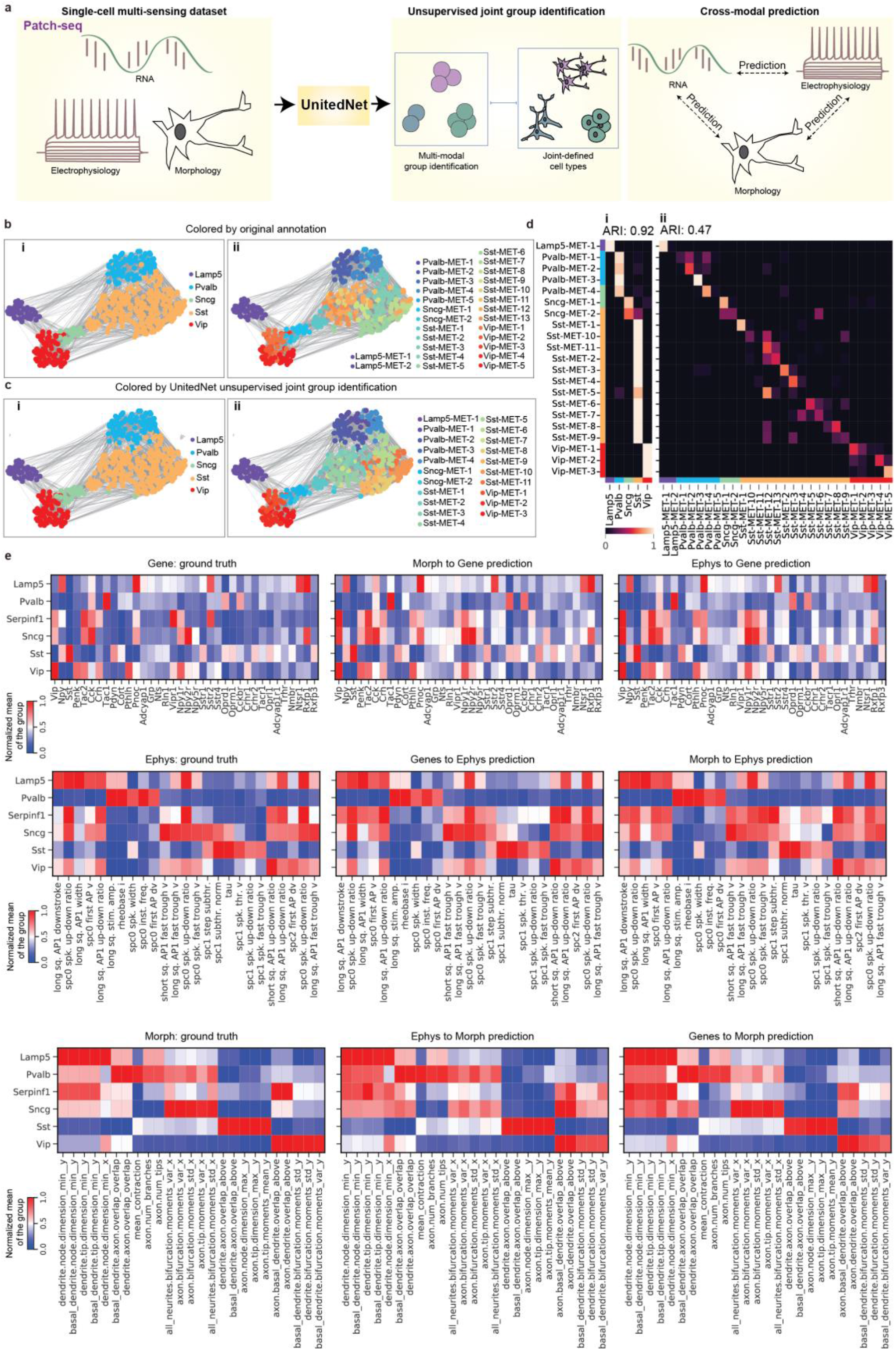
Multi-modal analyses of a multi-sensing dataset by UnitedNet. **a**, Schematics illustrating the unsupervised multi-modal group identification and cross-modal prediction of a Patch-seq dataset by UnitedNet. The patch-seq experiment simultaneously measures transcriptomics, electrophysiology, and morphology from the same cells to jointly define cell types. **b-c**, UMAP representations of the shared codes in the shared latent space that are color-coded by the joint morphology-electrophysiology-transcriptomics (MET)-types labeled by the reference (b) and identified by UnitedNet (c), where MET major cell type annotations are in i) and MET cell subtype annotations are in ii). **d**, Confusion matrices comparing joint major cell types and cell subtypes between the reference labels and UnitedNet-identified labels. **e**, Heatmap comparing cross-modal predicted gene expression, electrophysiological, and morphological features averaged over annotated cell types with the ground truth.

We first benchmarked the unsupervised group identification performance of UnitedNet by combining cell electrophysiological and morphological features to identify transcriptomic cell types^34^. Compared with other multi-modal cell typing methods, the performance of cell typing results by UnitedNet demonstrates substantial improvement in terms of cell type separability and identification accuracy (Extended Data Fig. 4). We then benchmarked the cross-modal prediction performance for electrophysiology-to-transcriptomics (E-T) by comparing UnitedNet with Coupled Autoencoder (CplAE), a deep neural network with an encoder-decoder structure^12,13^. UnitedNet achieved a better performance in all directions of cross-modal prediction than CplAE with different hyperparameters (Extended Data Fig. 5).

Next, we performed simultaneous unsupervised joint group identification analysis and cross-modal prediction on the morphological-electrophysiological-transcriptomic (MET) datasets. By directly fusing the three modalities together and assigning the label for each cell, UnitedNet identified the cell MET-types in an end-to-end manner with a high degree of congruence and a roughly diagonal correspondence between the major MET-types and subtle MET-types compared with the previously reported results (Fig. 3b-d). Notably, to the best of our knowledge, the current methods have only been applied to cross-modal prediction between two modalities. Through the cross-modal prediction analysis among all the three modalities of the Patch-seq dataset, UnitedNet enabled the prediction of individual measurements across three modalities with high fidelity (Fig. 3e, Extended Data Fig. 6).

### UnitedNet-enabled multi-modal analysis for multi-omics data

After the initial demonstration, we applied UnitedNet to analyze complex and large-scale datasets from multiple batches of samples. These large-scale multi-omics datasets are typically generated to annotate cell molecular types from different batches of samples. For example, the single-cell transposase-accessible chromatin sequencing (ATAC-seq) technique that combines gene expression and genome-wide DNA accessibility has been used to profile diverse types of immune cells^5^. Then, a key analytical challenge is to use the previous annotated multi-modality datasets as references to analyze the new measurement from a different batch of the same biological system. Meanwhile, another prevalent challenge is the batch effect, where the difference between cells is caused by the inter-sample variations. Because of this batch effect, it is difficult to use the labeled ATAC-seq dataset as a reference atlas to annotate other new datasets. We found that UnitedNet can address these challenges in a twofold manner. First, UnitedNet can achieve a supervised group identification task (termed annotation transfer) that automatically identifies cell types in new unlabeled test samples based on previously labeled training samples and simultaneously enable cross-modal prediction. In addition, we found that the supervised group identification combined with the alternating training can effectively reduce the batch effects among different biological samples. (Fig. 4a).

**Fig. 4.**
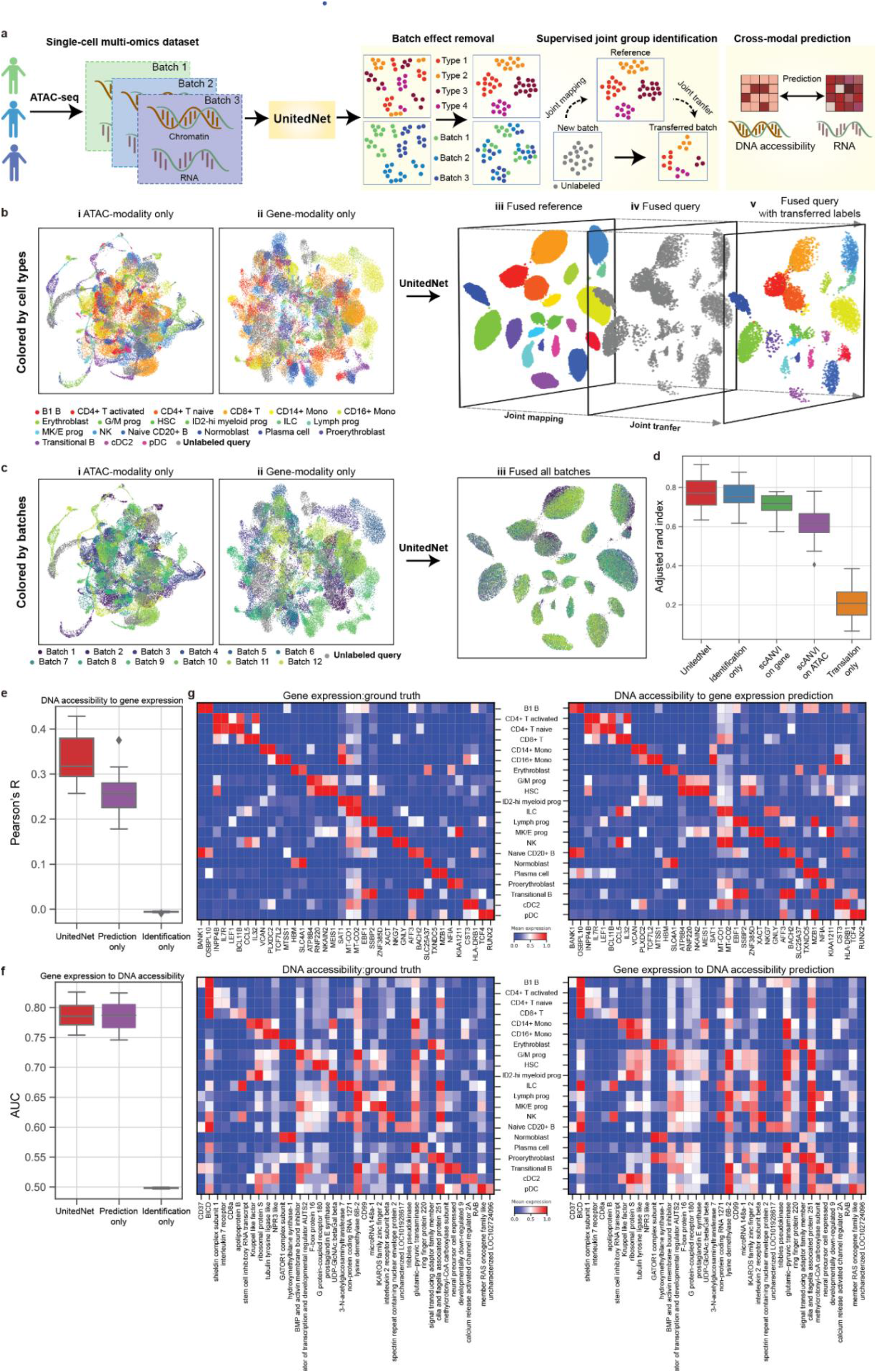
Automated multi-modal annotation transfer and cross-modal prediction of multi-omics datasets by UnitedNet. **a**, Schematics of an ATAC-seq data analyses pipeline by UnitedNet. ATAC-seq simultaneously measures RNA and chromatin accessibility in the same cells at the single-cell resolution. UnitedNet uses RNA and chromatin accessibility data along with their cell type annotation labels as inputs to train the network, simultaneously enabling batch-effect removal, supervised joint group identification, and cross-modal prediction. **b**, Supervised joint group identification enabled by UnitedNet. The UMAP latent space visualizations show DNA accessibility (i), RNA (ii), and shared latent codes of UnitedNet (iii-v) of the 13-batch ATAC-seq dataset. In this process, UnitedNet first fused the shared latent codes from two modalities and grouped the codes based on cell type annotations (iii). Then, it projected the unlabeled query batch to the learned shared latent space (iv), transferring the label of the reference to the shared latent codes of the unlabeled query batch to complete the annotation transfer (v). The latent codes are colored by cell-type annotations. **c**, Joint batch effect removal enabled by UnitedNet. The UMAP latent space visualizations show DNA accessibility (i), RNA (ii), and shared latent codes (iii) from a 13-batch ATAC-seq dataset. The latent codes are colored by different batches. **d**, Quantitative joint group identification results of UnitedNet and other ablation and benchmarking methods (*scANVI*^35^). *scANVI* is used for single-modality annotation transfer based on the ATAC or the gene expression modality. **e-f**, Quantitative prediction results of UnitedNet and other ablation studies on the ATAC-seq dataset. The prediction from DNA accessibility to RNA (e) and the prediction from RNA to DNA accessibility (f) are evaluated by Pearson’s *r* and the area under the curve (AUC), respectively The boxes depict mean+/− standard error (S.E.) over leave-one-out cross-validations, in which 12 of 13 batches are used as training data and the rest one batch as testing data. **g**, Heatmap comparing cross-modal predicted gene expression and DNA accessibility features with the ground truth. The values are averaged and min-max scaled over annotated cell types.

We applied UnitedNet on ATAC-seq datasets measured from the bone marrow mononuclear cells (BMMCs) in 13 batches of different tissue sites and donors^7^. In these datasets, 21 cell types were previously identified and annotated (Fig. 4b i-ii and Fig. 4c i-ii). We trained UnitedNet by using 12 batches of data. The corresponding cell type annotations were used as the labels of training samples. Then, we tested the performance on the 13^th^ batch. UnitedNet successfully (i) integrated 12-batch reference datasets and learned 21 annotated cell types, (ii) simultaneously mapped the unlabeled test dataset (the 13^th^ batch) into the same 21 separated clusters, and (iii) removed the batch effect (Fig. 4b iii-v and Fig. 4c iii). Next, we validated whether multi-task learning can still achieve better performance in a supervised setting using ablation studies. Compared with a *state-of-the-art* annotation transfer method^35^ and ablated UnitedNet in which only one task is involved, UnitedNet trained with supervised group identification and cross-modal prediction showed the highest accuracy in both tasks (Fig. 4d-f). Finally, we asked whether the cross-modal prediction by UnitedNet could reconstruct cell-type-specific feature patterns. We compared the gene expression and DNA accessibility patterns across cell types between the ground truth and predicted results. The high similarity in gene expression-to-DNA accessibility prediction (Pearson’s *r* = 0.96 ±0.03) and DNA accessibility-to-gene expression (Pearson’s *r* = 0.77 ± 0.10) demonstrated the high performance of the cross-modal prediction task (Fig. 4g). Collectively, UnitedNet showed enhanced performance for ATAC-seq data analysis by combining annotation transfer and cross-modal prediction tasks for multi-task learning.

### UnitedNet-enabled multi-modal analysis for spatial-omics data

Finally, we extended UnitedNet to spatial-omics datasets, which are generated by measuring spatially resolved multi-omics information in intact tissues^36–38^. Traditionally, spatial-omics datasets are not considered multi-modality datasets. UnitedNet, however, can take spatial information as one modality for multi-modal analysis, thereby combining spatial and multi-omics information to identify biologically meaningful groups and enable cross-modal prediction. Specifically, we created an independent modality by extracting neighborhood gene expression information to represent the niche identity (See Methods, Fig. 5a).

**Fig. 5.**
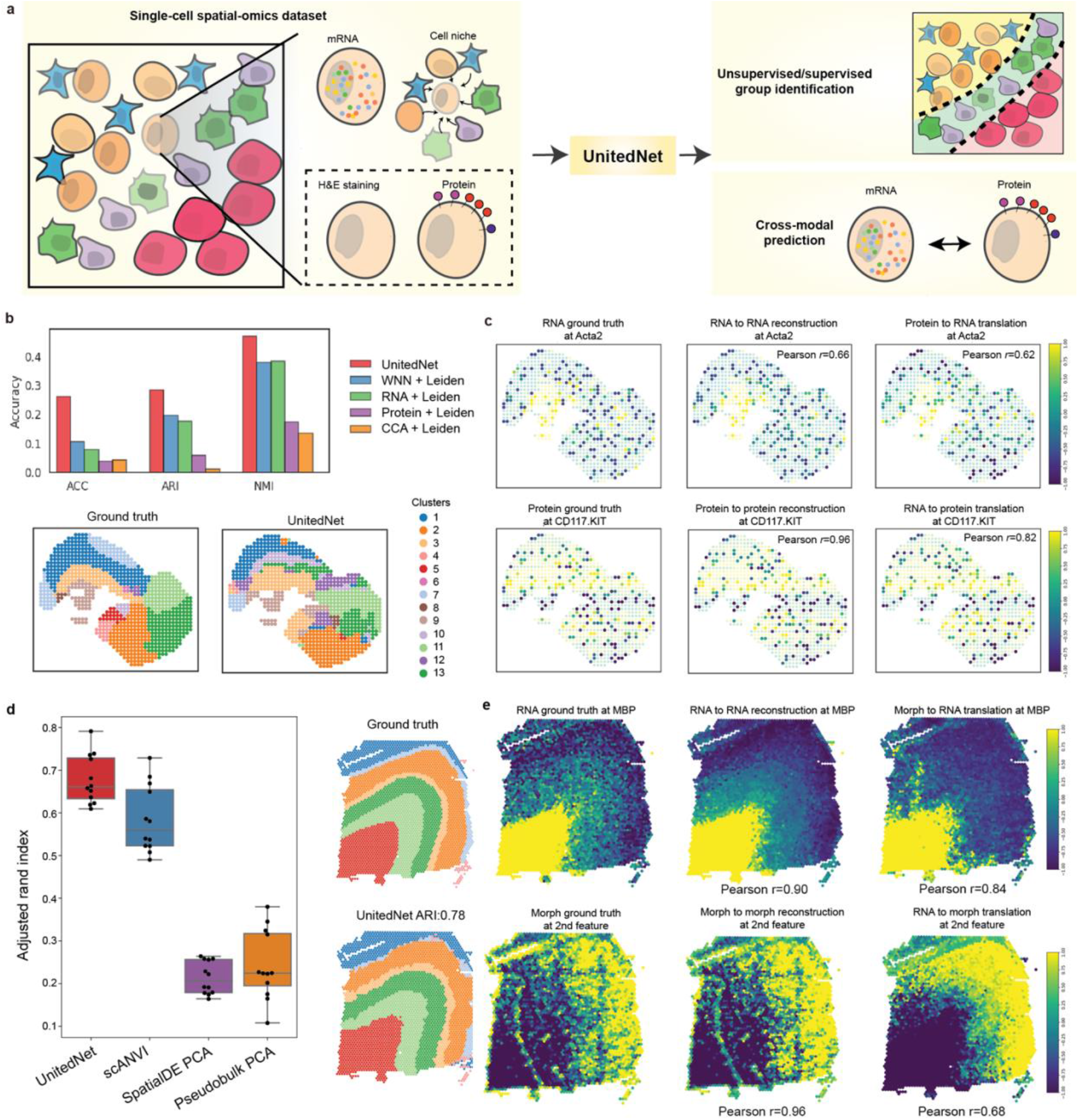
Automated unsupervised group identification, multi-modal annotation transfer, and cross-modal prediction of spatial-omics datasets by UnitedNet. **a**, Schematics of spatial-omics data analyses pipeline by UnitedNet. Spatial omics simultaneously measure spatially resolved multi-omics data in intact tissue networks. UnitedNet extracts the cell neighborhood information as an additional modality together with other modalities for unsupervised/supervised group identification and cross-modal prediction. **b**, The performance comparison between UnitedNet and other methods for tissue region group identification on the DBiT-seq embryo dataset. **c**, Results of cross-modal prediction between representative genes and proteins. **d**, Box and dot plots comparing the performance of supervised group identification for tissue region identification on the DLPFC dataset among UnitedNet, *SpatialDE PCA*, and *Pseudobulk PCA* from the previous study, and *scANVI*. *scANVI* is used for single-modality annotation transfer based on the gene expression modality. Each dot represents the performance with the combination of 11 batches as training reference and 1 remaining batch as the test dataset. The boxes represent mean+/− standard error (S.E.). **e**, Prediction results of representative genes, and morphological features from the DLPFC dataset.

We first applied UnitedNet on a single-batch DBiT-seq embryo dataset that simultaneously mapped the whole transcriptome and 22 proteins on an embryonic tissue^30^. Specifically, we generated the weighted average of RNA expression in the niche that encodes spatial information as the third modality for analysis. UnitedNet then combined gene expression, protein, and niche modality for unsupervised joint identification of tissue regions and cross-modal prediction between gene expression and proteins. We benchmarked the accuracy of the tissue region identification by considering the anatomic annotation of the tissue region from the original report as ground truth. UnitedNet achieved a higher unsupervised group identification accuracy compared with *state-of-the-art* methods (Fig. 5b). In addition, UnitedNet enabled the spatially resolved cross-modal prediction between gene and protein expressions (gene expression-to-protein, Pearson’s *r* = 0.451 ± 0.08, protein-to-gene expression, Pearson’s *r* = 0.185 ± 0.061; Fig. 5c).

Next, we applied UnitedNet to an annotated multi-batch spatial-omics dataset for simultaneous supervised joint group identification and cross-modal prediction. We used a human dorsolateral prefrontal cortex (DLPFC) dataset that spatially maps the gene expression and H&E staining on the 6-layered DLPFC brain slices from 12 batches^29^. Similarly, we used the gene expression, H&E staining-based morphological features, and niche modality as inputs to UnitedNet. UnitedNet can successfully annotate unseen DLPFC slices (Extended Data Fig. 7) and achieve higher accuracy than the other benchmarking methods and ablated UnitedNet in which we removed the niche modality (Fig. 5d and Extended Data Fig. 8). Furthermore, UnitedNet can enable the cross-modal prediction between the gene expression and the H&E morphological features (gene expression-to-H&E features, Pearson’s *r* =0.120±0.022 and H&E features-to-gene expression, Pearson’s *r* = 0.273±0.003; Fig. 5e).

In brief, UnitedNet offers a new direction that extracts the spatial information as an input modality, to enable supervised and unsupervised group identification and cross-modal prediction for spatial-omics data.

## Discussion

We have demonstrated that UnitedNet can integrate the joint group identification and cross-modal prediction tasks through multi-task learning for single-cell multi-modality data analysis. Through extensive ablation and benchmarking studies, we have validated that multi-task learning can achieve better performance than single-task learning, single-modality analysis, and other *state-of-the-art* methods in both unsupervised and supervised settings. In addition, UnitedNet may be applicable to various single-cell multi-modality biological datasets, including, but not limited to, multi-modality simulation data, multi-sensing data, multi-omics data, and spatial omics data. The success of UnitedNet in analyzing multi-modality biological data will significantly expand our ability to chart and predict cell states via combined multi-modality information in heterogeneous biological systems. We envision that multi-task learning can be used for data-driven scientific discoveries such as neuron subtype clustering, stem cell maturation state identification, and spatially resolved multi-modal brain mapping.

Finally, although the UnitedNet performed well for different multi-modality datasets, a number of improvements can be introduced, such as automatic searching for the network configuration (*e.g*., the numbers of neuron nodes used in each layer) and optimal hyperparameter (*e.g*., the learning rate that controls the optimization level in each training iteration) for deep neural networks^39^.

Moreover, there are some remaining questions such as why multi-task learning can improve multi-modal data analysis, whether the network can integrate more tasks *(e.g.*, the identification of cross-modal feature relevance) with group identification and cross-modal prediction tasks, and how to leverage different tasks to design customized algorithms for particular datasets^18,19^. We envision that future mechanistic studies, including ablation experiments and theoretical analysis, could further help us address these questions.

## Methods

### UnitedNet

UnitedNet features both cross-modality prediction and joint group identification. The above two tasks are accomplished based on learned joint low-dimensional representations of different modalities, which contain the essential information of the cells (Fig. 1b). Suppose that there are *V* different modalities. Let *n* denote the number of cells and *p*^(*ν*)^ denote the number of features in the *ν^th^* modality. Let 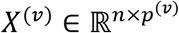 (*ν* = 1, …,*V*) denote the data from modality *ν* where its *i^th^* row 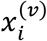 corresponds to the cell *i*. For the prediction from modality *ν*_1_ to *ν*_2_, UnitedNet predicts 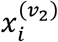 from the latent code (low-dimensional representation) obtained from 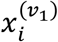. For group identification, UnitedNet first fuses the latent codes of 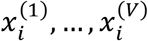. Next, based on the shared latent codes, it returns a group index *k*_i_ ∈ {1, …, *K*}.

UnitedNet consists of encoders, decoders, discriminators, and a group identification module. It is trained based on a within-modality prediction loss, a cross-modality prediction loss, a generator loss, a discriminator loss, a contrastive loss, a clustering loss (for unsupervised group identification), and a classification loss (for supervised group identification). The details about the components and losses of UnitedNet are as follows.

### Encoders

For each modality *ν* = 1, …, *V*, UnitedNet has one encoder *Enc*^(*ν*)^(·) that maps the features of each cell *i* (*i* = 1, …, *n*) to a modality-specific latent code 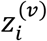 containing the most essential information of the data:

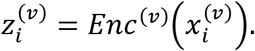

The low-dimensional representations from different modalities are required to have the same number of components.

### Decoders

The decoder *Dec*^(*ν*)^(·) takes modality-specific latent codes as the input and maps them to the features of modality *ν*. We denote the cross-modality predicted features from modality *ν*_1_ to *ν*_2_ (where *ν*_1_ ≠ *ν*_2_) by

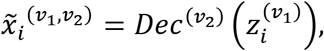

and the within-modality predicted features of modality *ν* by

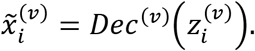

### Discriminators

Discriminators assist the training of the generator that consists of the encoders and decoders. The discriminator *Dis*^*ν*^(·) of modality *ν* takes either the within-modality predicted features 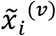 or original features 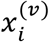 as the input and outputs a binary classification result, aiming to distinguish between 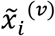 and 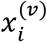. The encoders and decoders improve their performance by increasing the error rate of the discriminators.

### Joint group identification module

Denote the number of groups to be *K*. The group identification module takes the modality-specific codes from all modalities 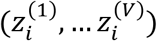 as the input and assigns it to one of the *K* groups. It first fuses the data:

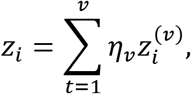

where *η*_1_ …, *η_V_* are positive trainable weights with 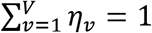 *η*_*ν*_ = 1. Next, the fused representation *z_i_* is passed through a fully connected layer:

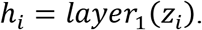

Then, to obtain the soft assignment of the group index, the intermediate output *h_i_* is processed by another fully connected layer:

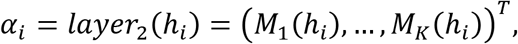

where

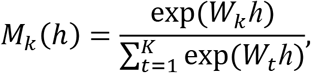

and *W_k_* (*k* = 1, …, *K*) is model coefficients. The group identification module assigns the group index 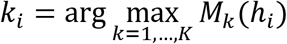 to cell *i*.

### Clustering loss

The clustering loss consists of three components. The first two components are adopted from the Deep Divergence-based Clustering (DDC)^41^. They ensure that the obtained groups are separable and compact. The third component is based on the self entropy^42^. It avoids trivial solutions where most of the cells are assigned to a small portion of the total number of groups.

#### Component 1

Component 1 reduces the two-by-two correlations between the cluster probability assignments (soft assignments) of different groups. It increases the group separability since those correlations are positively related to the similarity between different groups. Define the matrix 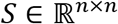 with matrix element 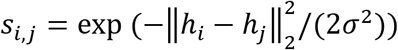, where *i* = 1, …, *n*, *j* = 1, …, *n*, and *σ* is a hyperparameter. It measures the similarity between different cells. Denote (*M_k_*(*h*_1_), …, *M_k_*(*h_n_*)) by 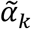, which is the soft assignments of for the group *k* for cells 1 to *n*. Then, the component 1 is calculated by

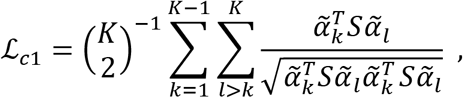

where 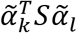 is an estimate of 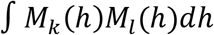, and it is minimized when *M_k_*(*h*) is orthogonal to *M_l_*(*h*). Accordingly, 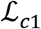 requires the values *M_k_*(·) and *M_l_*(·) (*k* ≠ *l*) to have low correlation.

#### Component 2

The component 2 pushes the soft assignment values of different groups to distinct corners of the simplexes in 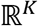, which also increases the group separability. Let 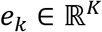 denotes a vector with its *k^th^* element be one and other elements be zero. Therefore, *e_k_* (*k* = 1, …, *K*) is the *k^th^* corner of the simplex. Recall that *α*_*i*_ is the output from *layer*_2_ in the group identification module. Let 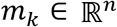 with its *i^th^* element be 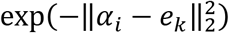, which measures the distance between the soft assignments *α_i_* and simplex corner *e_k_*. The component 2 is defined by

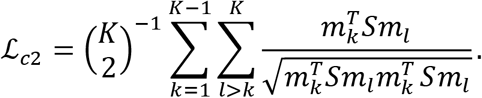

It enforces the orthogonality between 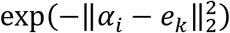 and 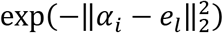. Therefore, the soft assignment output (*M*_1_(*h*), …, *M*_*K*_(*h*))^*T*^ will tend to get close to one simplex corner (e.g., *e_k_*) instead of approaching multiple of them (e.g., both *e_k_* and *e_l_*) at the same time. Consequently, the low-dimensional representation in the same group will be compact and those from different groups will be separated.

#### Component 3

Component 3 aims to avoid the trivial solution where most of the cells are assigned to a small proportion of the total groups. Let 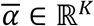 denote the averaged *α_i_* for *i* = 1, …, *N*, with its *k^th^* element denoted by 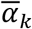 (*k* = 1, …, *K*). The third component is based on the entropy of 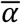:

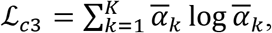

which aims to assign each group index with equal probability (*i.e.*, 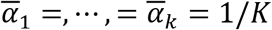). Therefore, it regularizes the groups by avoiding the assignment to be a few subsets of total clusters.

### Prediction loss

The within-modality prediction loss is defined by

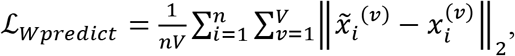

which measures the distance between the within-modality predicted features 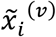 and the original features 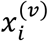. The cross-modality prediction loss is defined by

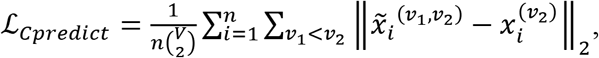

which measures the distance between the cross-modality predicted features from modality *ν*_1_ to *ν*_2_ and the original features from modality *v_2_*. When the low-dimensional representation captures the essential information of the data, both 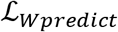 and 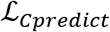 are expected to be small.

### Generator loss and discriminator loss

The generators and discriminators are trained by the least-squares loss^25^. We assign the within-modality predicted features 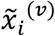 with label one and the original features 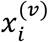 with label zero. The generator loss is defined by

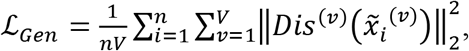

which is minimized when 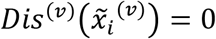 for *i* = 1, …, *n* and *ν* = 1, …, *V*, namely, when the discriminator incorrectly classifies all the within-modality predicted features to zero. The discriminator loss is defined by

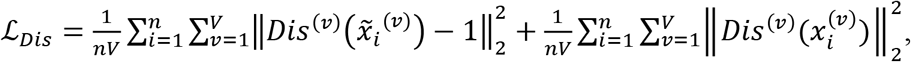

which aims to make the discriminator classify the within-modality predicted features to one and original features to zero. This set of least-squares losses improves the quality of the trained generator, since it matches the essential goal of the generator, which is generating feature data with a distribution similar to that of the original features.

### Contrastive loss

We apply the contrastive loss^31,43^ to align the latent codes from different modalities. Define the cosine similarity by

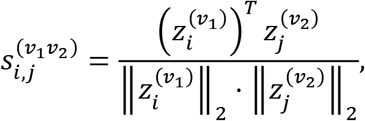

where *i*, *j* = 1, …, *n*. It is maximized when *z_i_*^(*ν*_1_)^ and *z_j_*^(*ν*_2_)^ are parallel. Let

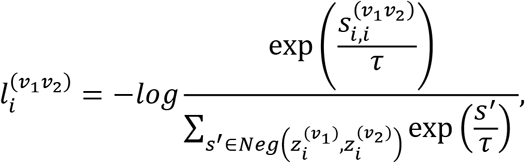

where 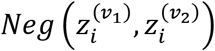 is obtained by sampling a fixed number of elements from the set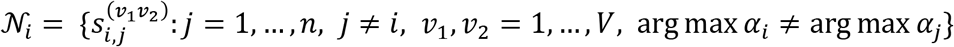, and *τ* is a hyperparameter. The term 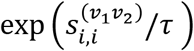 aligns the modality-specific codes. For the denominator, the set 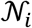 contains all the cosine similarities between the latent codes of cell *i* and those from the other cells that are grouped into different groups. The contrastive loss is defined

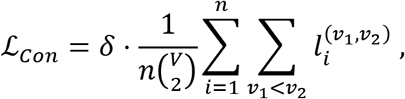

by where *δ* is a hyperparameter.

### Classification loss

Define vector *b_i_*, with its fcth element be one and the other *K* – 1 elements be zero, where *k* is the observed class label of cell *i*. We assess the classification accuracy of the group identification module by the cross-entropy:

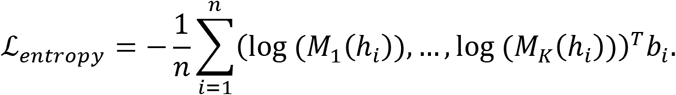

### Training procedure

We first present the procedure where UnitedNet is trained without cell labels for cross-modality prediction and unsupervised group identification. It is trained iteratively with two steps: *group identification update step* and *prediction update step*. In the *group identification update step*, the encoder outputs the modality-specific codes and feeds them to the group identification module. Then, the group identification module fuses the modality-specific codes as shared latent codes and obtains the *K*-dimensional soft cluster assignments. Next, the encoders and clustering module are updated by the group identification:

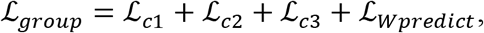

where the within-modality prediction loss 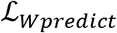 captures the intrinsic cell type information from the data. Without 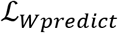, the unsupervised group identification module may undesirably generate clusters based on the noise from the data. In the *prediction update step*, the encoders output the low-dimensional representations from different modalities, feed them to the decoders, and obtain the cross-modality predicted features 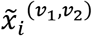 together with within-modality predicted features 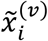. Then, the within-modality predicted features are fed into the discriminators. The discriminator is updated by the discriminator loss 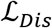. Next, the encoder and decoder are updated by the sum of the within-modality prediction loss, cross-modality prediction loss, generator loss, and contrastive loss:

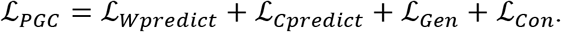

The above two training steps are summarized in Algorithms 1 and 2, respectively.

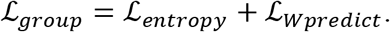

To train UnitedNet for supervised classification, we modify the group identification loss by:

#### Algorithm 1 Group identification update step

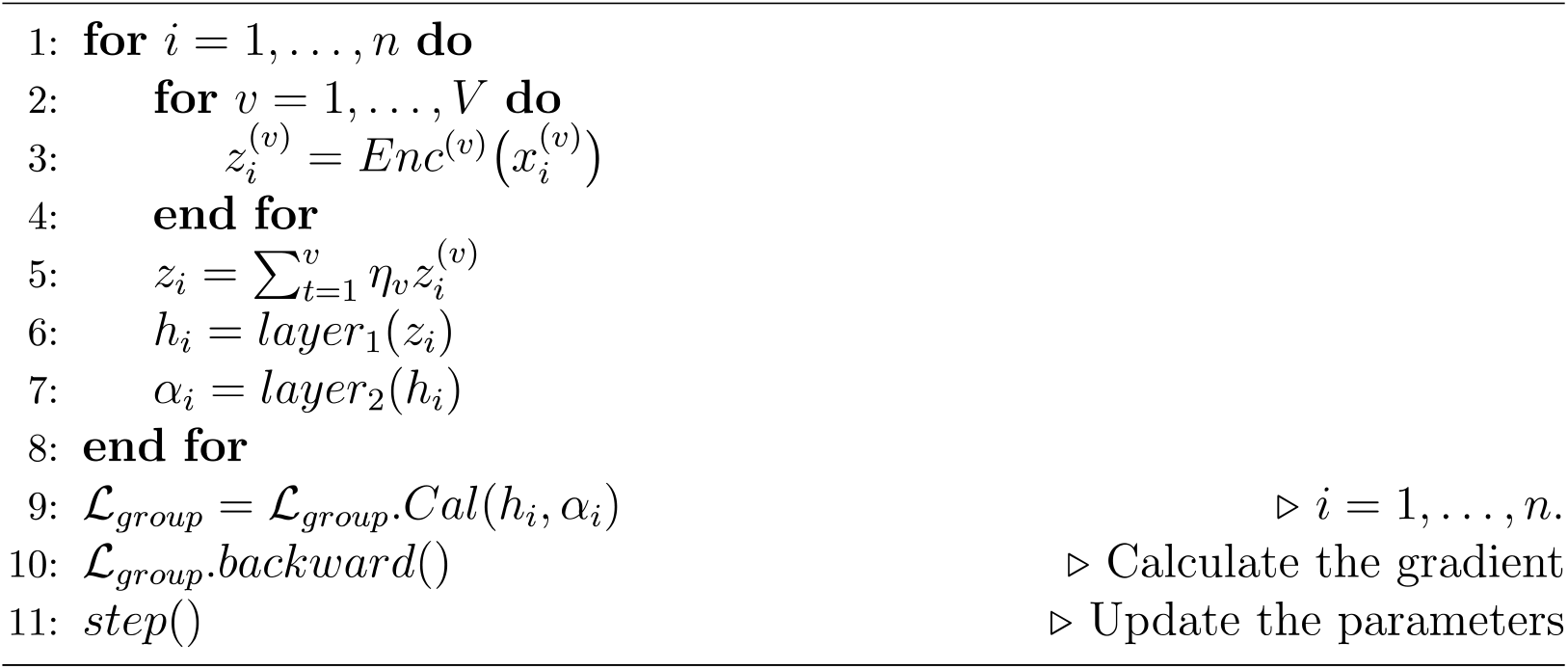

#### Algorithm 2 Prediction update step

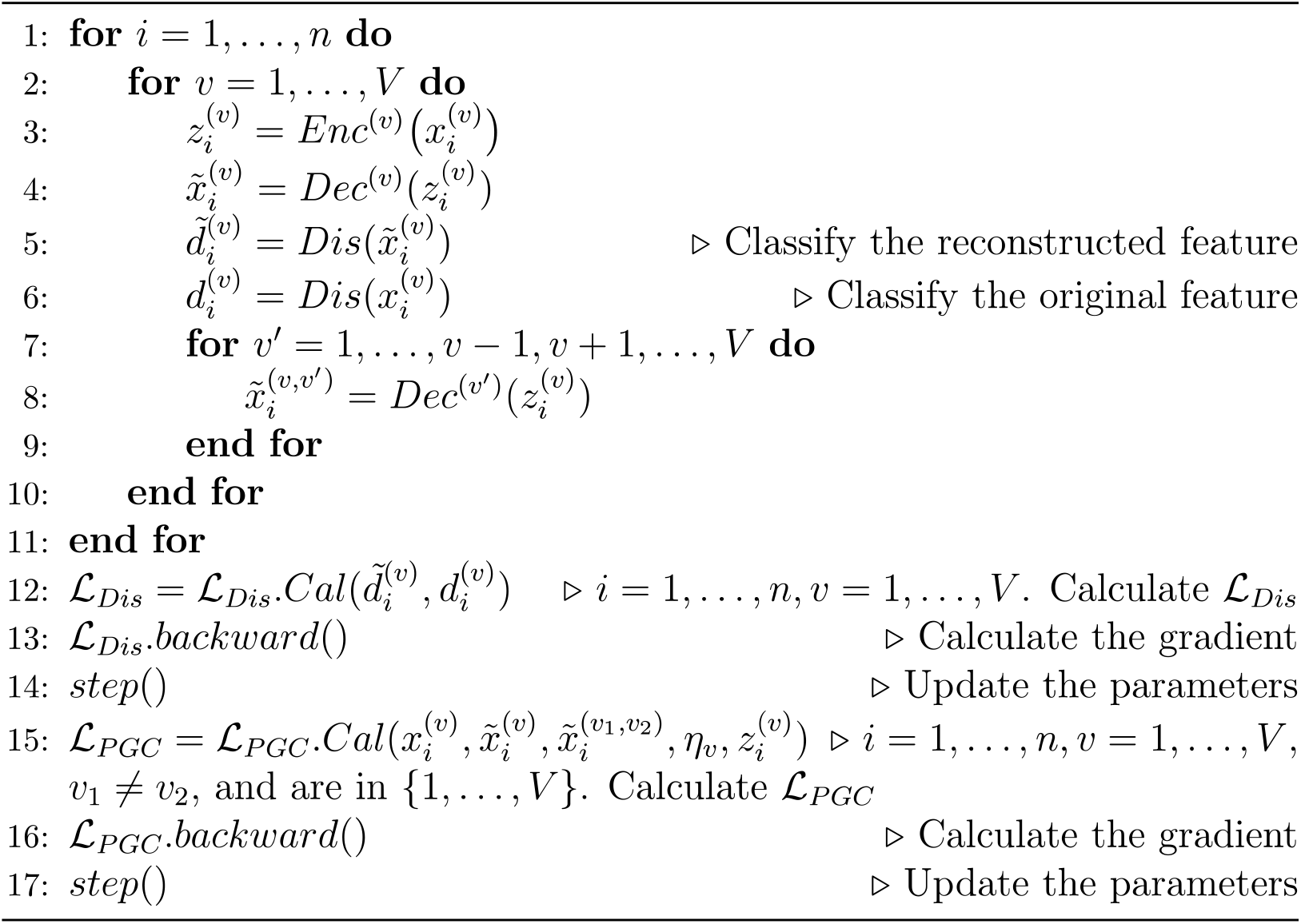

### Evaluation metrics

The clustering (unsupervised group identification) performance is evaluated by the adjusted rand

index and normalized mutual information. The prediction performance is evaluated by the coefficient of determination, Pearson’s correlation, and area under the ROC curve. All metrics are calculated by *scikit-learn^44^*. The details about these metrics are as follows.

### Adjusted rand index

The adjusted rand index compares the clusters obtained from the model with the one from the cell type labels. Let *a_k_a__* (*k_a_* = 1, …, *K*) denote the number of observations in the *k*th cluster from the model and *b_k_b__* (*k_b_* = 1, …, *K*) denote the number of observations in the *k*th cluster from the cell type labels. Let *n_k_a__,*k_b_** (*k_a_* = 1, …, *K*, *k_b_* = 1,…, *K*) denote the number of observations in both the *k_a_* th cluster from the model and *k_b_*th cluster from the cell type labels. The adjusted rand index is calculated by

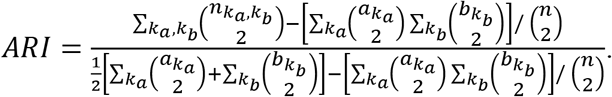

*ARI* takes a value between zero and one. A value close to one indicates that the clustering result from the model is close to the one from the cell type labels.

### Normalized mutual information

The normalized mutual information is calculated by

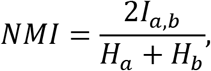

Where 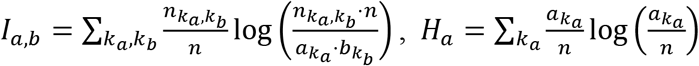, and 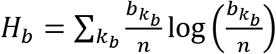 *NMI* takes a value between zero and one, and it gets close to one when the clustering result from the model is close to the one from the cell type labels.

### Coefficient of determination

Let *y* and 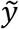 denote the observed data and predicted data, respectively. Define 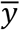 to be the vector with the same length as *y* and 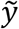 and each of its elements equals the averaged value of *y*. The coefficient of determination is calculated by

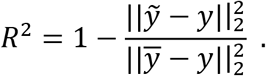

It compares the MSE of the prediction 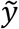 from the model with the MSE of the baseline that takes the constant value 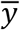 as the prediction. It takes a value from (−∞, 1] and equals one when 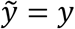.

### Pearson’s Correlation

Let 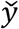 denote a vector with the same length as *y* and each of its elements equals the averaged value of *y*. The Pearson’s correlation is calculated by

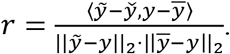

It takes a value in [−1,1] and equals one or negative one when the prediction has positive or negative linear relationship with the observed data, respectively. It equals zero when the prediction and observed data have no linear relationship.

### Area under the curve (AUC)

For modeling the ATAC-seq dataset, we have binarized the chromatin accessibility data. Thus, to evaluate the prediction for those data, we adopt the area under the ROC curve, which is already applied in the previous study^13^. Let *n*_0_ and *n*_1_ denote the number of zeros and ones from the observed data, respectively. Let *p*_0,*i*_ (*i* = 1, …, *n*_0_) and *p*_1,*j*_ (*j* = 1, …, *n*_1_) denote the observations from the two groups, respectively. The area under the ROC curve is calculated by:

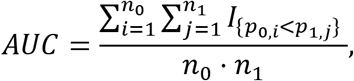

where

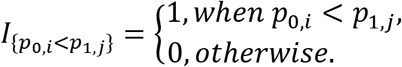

*AUC* takes values between zero and one, and one corresponds to a perfect prediction.

### Dataset used for multi-task learning in UnitedNet

The input of UnitedNet can be any single-cell multi-modality dataset. We conclude that four major categories of such datasets include multi-modal simulation datasets, multi-modal sensing datasets, multi-omics datasets, and spatial omics datasets. The experimental details are specified in the following sections.

### Multi-modal datasets used for the demonstration of UnitedNet

#### Dyngen simulated dataset

We use *Dyngen^26^* to simulate the four-modality dataset. Specifically, we generate 500 cells with simulated DNA, pre-mRNA, mRNA, and protein modalities, each modality containing 100 dimensions of features. Meanwhile, the ground truth cell-type annotations are generated along with the dataset. For the parameter of the Dyngen simulator, we use the default setting of a linear backbone model in the tutorial of Dyngen with the functions including *backbone_linear, initialize_model*, and *generate_dataset*.

#### MUSE simulated dataset

We apply the simulator in *MUSE^27^* to simulate two-modalities input to assess the robustness of UnitedNet with one low-quality. We simulate 11 two-modality datasets with 1,000 cells and 10 cell types. Each modality contains 500 modality-specific features. For each of the 11 datasets, one of the modalities is simulated with a controllable dropout coefficient. We use 0.01, 0.1, 0.2, 0.3, 0.4, 0.5, 0.6, 0.7, 0.8, 0.9, and 1 as different dropout coefficients when benchmarking with other methods.

#### MNIST multi-modal dataset

We use the custom-built MNIST^33^ as the four-modality input for UnitedNet to further validate the performance of UnitedNet. MNIST is a validation dataset with 70,000 images of hand-written digits in which 60,000 images are used as the training dataset and the remaining 10,000 images are used as the test dataset. We generate a four-modality dataset including the original, edge-transformed, invert-transformed versions of MNIST as well as one pure noise modality. We use *OpenCV 4.5.5* to conduct the edge transformation and invert transformation.

#### Patch-seq GABAergic neuron dataset

We use a Patch-seq dataset that simultaneously characterizes the morphological (M), electrophysiological (E), and transcriptomic (T) features obtained from GABAergic interneurons in the mouse visual cortex^28^. We use the same dataset after quality control of previous research, in which 3,316 neurons remain for E-T analysis and 438 neurons remain for M-E-T analysis^12,28^. We standardize the input matrices of each modality to make the mean value and the standard deviation of all features in each cell to be 0 and 1, respectively.

#### ATAC-seq BMMC dataset

We use an ATAC-seq dataset that simultaneously combines gene expression and genome-wide DNA accessibility obtained in BMMC tissue from 10 donors and 4 tissue sites^21^. In addition to the quality control in the previous study, we use the standard preprocessing procedure for the ATAC-seq BMMC datasets. For the preprocessing of the gene expression modality, we use median normalization and the log1p transform and standardization and select the top 4,000 most variable genes through *Scanpy*^45^. For the preprocessing of the DNA accessibility modality, we binarize the data by replacing all nonzero values with a value of 1 and select the top 13,634 most variable DNA accessibility features through *Scanpy*. We use the *ChIPseeker 3.14*^46^ to annotate the DNA-accessibility peaks.

### UnitedNet on spatial omics datasets

#### Generating the niche expression modality

Using the measured expression of RNAs of cells or spots, we incorporate the spatial information of each cell or spot and generate a weighted average expression of RNAs. With two-dimensional spatial coordinates 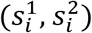 and modality *ν* with its *i*^th^ row 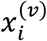 corresponding to the cell/spot *i*, we compute the niche modality for modality *ν* that is denoted by *x*^(*ν niche*)^ with (*ν* = 1, …, *V*). For cell/spot *i*, we compute 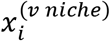 by:

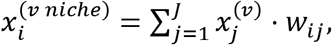

where *j* ∈ {1, …,*J*} denotes cells/spots that belong to the K-nearest neighbors of cell/spot *i*, and *w_ij_* is calculated by:

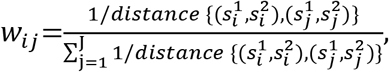

where *distance*{·} denotes the Euclidean between two vectors.

#### UnitedNet on DBiT-seq embryo dataset

We use the DBiT-seq embryo dataset^30^, where the following four modalities of 936 spots in DBiT-seq are taken: mRNA expression, protein expression, and niche mRNA expression. For modality of mRNA expression, we normalize the raw count matrix using function *scanpy.pp.normalize_total* from *scanpy* and select the top 568 differentially expressed genes. For modality of protein expression, we normalize the raw count matrix and used 22 kinds of proteins. The niche modalities are generated based on normalized mRNA expression. For the first task of tissue region characterization, ground truth tissue region labels are extracted from the original research^30^ which is the anatomic annotation of major tissue regions based on the H&E image. We compare clustering results from UnitedNet with those from WNN combined with *Leiden* clustering, *Leiden* clustering on the single modality of mRNA expression, *Leiden* clustering on the single modality of protein expression, and canonical correlation analysis combined with *Leiden* clustering. We validate their performance by adjusted rand index. For the parallel task of translation across modalities, although three modalities are used as input to the UnitedNet model, we focus on prediction and translation between the first and second modalities: mRNA expression and protein expression. Since there is one batch in DBiT-seq public datasets, we split the total 936 spots in the DBiT-seq embryo dataset into the training dataset (80%, 748 spots) and test dataset (20%, 188 spots) for the prediction and translation task.

#### UnitedNet on DLPFC dataset

We use the human adult dorsolateral prefrontal cortex (DLPFC) datasets with 12 batches^29^. We use the following three modalities of 936 spots in DBiT-seq: mRNA expression, morphological features extracted from the H&E staining image, and niche mRNA. For modality of mRNA expression, we normalize the raw count matrix and selected the top 2365 differentially expressed genes. We use a pre-trained convolutional neural network^47^ to extract morphological features from the H&E staining image implemented by *stLearn^48^*. A 50-dimensional morphological feature is used as the second modality for each spot. For the supervised group identification task, we use 11 batches with their tissue region annotations to train a UnitedNet model. Then we apply the trained model to the remaining batch to identify tissue region annotation. We compare the identification performance of UnitedNet with *SpatialDE PCA* and *pseudobulk PCA* from the original DLPFC paper^29^.

## Data availability

The Dygen simulation data can be reproduced by the simulator in https://github.com/dynverse/dyngen. The MUSE simulation data can be reproduced by the simulator in https://github.com/AltschulerWu-Lab/MUSE. The original modality of MNIST data was downloaded from http://yann.lecun.com/exdb/mnist. The Patch-seq GABAergic neuron dataset was downloaded from https://github.com/AllenInstitute/coupledAE-patchseq and https://portal.brain-map.org. The ATAC-seq BMMC dataset was downloaded from https://openproblems.bio/neurips. The DBiT-dataset was downloaded from: https://www.ncbi.nlm.nih.gov/geo/query/acc.cgi?acc=GSE137986. The DLPFC dataset was downloaded from https://doi.org/10.18112/openneuro.ds002076.v1.0.1.

## Code availability

Source code and demonstration code will be made available at https://github.com/LiuLab-Bioelectronics-Harvard/UnitedNet upon publication. A Python package of UnitedNet will be released and can be installed through ‘pip install unitednet’ upon publication.

## Acknowledgments

We thank Jane Salant for her helpful comments on the manuscript. J.L., J.D., and N.L. acknowledge the support from the NSF ECCS-2038603. J.L. acknowledges the support from NIH/NIDDK 1DP1DK130673 and William F. Milton Fund. Y.H. acknowledges the support from the James Mills Peirce Fellowship from the Graduate School of Arts and Sciences of Harvard University.

## Author contributions

J.L., J.D., and X.T. conceived the idea and designed the research. X.T. designed and developed the model. J.Z. provided critical discussions during the development. X.T. and Y.H conducted analyses on simulated and biological datasets. X.T. packaged the model as a pipeline. All authors prepared figures and wrote the manuscript. J.L. and J.D. supervised the study.

## Additional information

Correspondence and requests for materials should be addressed to J.L. and J.D.

---

**Extended Data Fig. 1.**
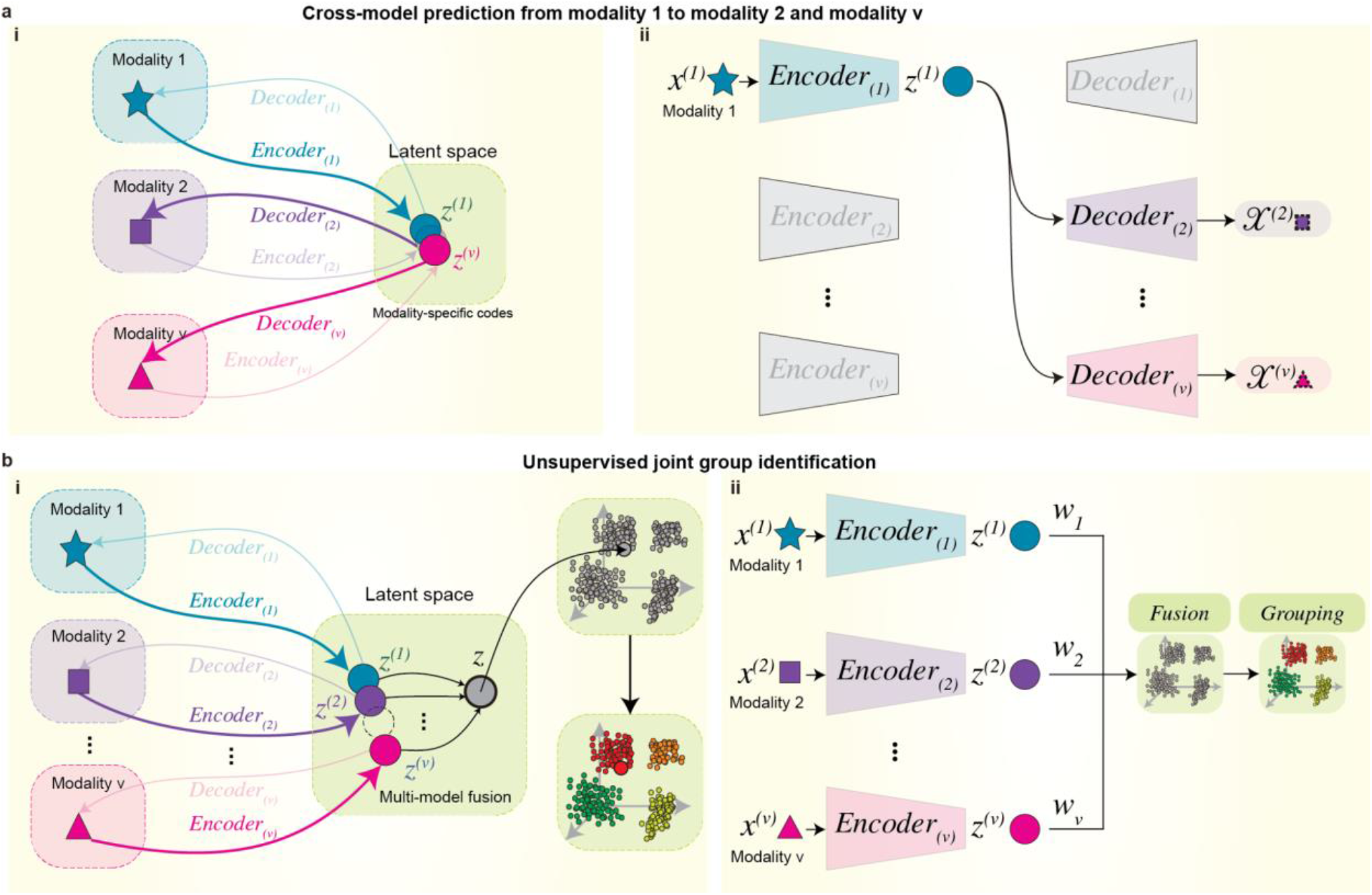
Tasks in UnitedNet and the multi-task learning procedure. **a**, An example of a cross-modal prediction task. **i**, The modality 1 is encoded to the latent space as modality-specific codes by encoder 1. The modality-specific codes are then decoded to the space of modality 2 and modality v through decoder 2 and decoder v. (ii), The network structure used for the cross-modal prediction task in (i). **b**, An example of a joint cell typing task. (i) Modality 1, modality 2, and other modalities are encoded to the latent space as modality-specific codes through corresponding encoders. The modality-specific latent codes are then fused as a shared latent code. The shared latent code is then used for unsupervised/supervised group identification through the grouping module. (ii) The network structure used for the joint group identification task in (i).

**Extended Data Fig. 2.**
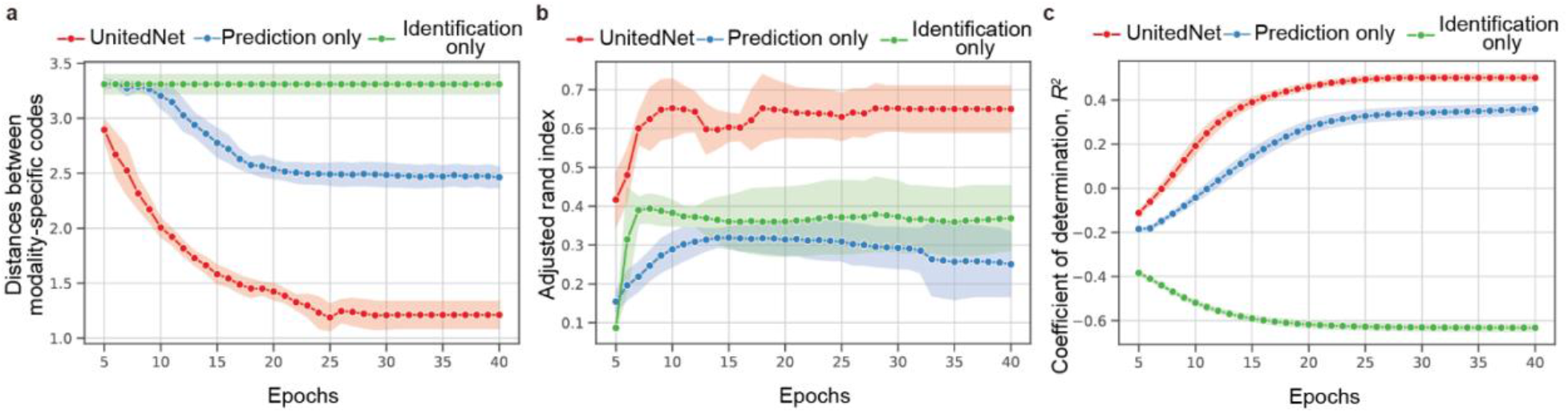
Ablation analysis of UnitedNet on a simulated Dyngen dataset. **a**, The line plot shows the change of the average distance of modality-specific codes from the latent space of UnitedNet with different numbers of training epochs. UnitedNet trained with multi-task learning shows the best performance of alignment among other ablations. **b**, The line plot shows the change in the performance of joint group identification that is measured by the adjusted rand index with different numbers of training epochs. UnitedNet achieved the highest performance among other ablations. **c**, The line plot shows the change in the performance of cross-modal prediction as measured by the coefficient of determination, *R*^2^, with different numbers of training epochs. UnitedNet achieves the highest performance among other ablations. Error bars show mean ± s.d. over 5-fold cross-validation for panels a-c.

**Extended Data Fig. 3.**
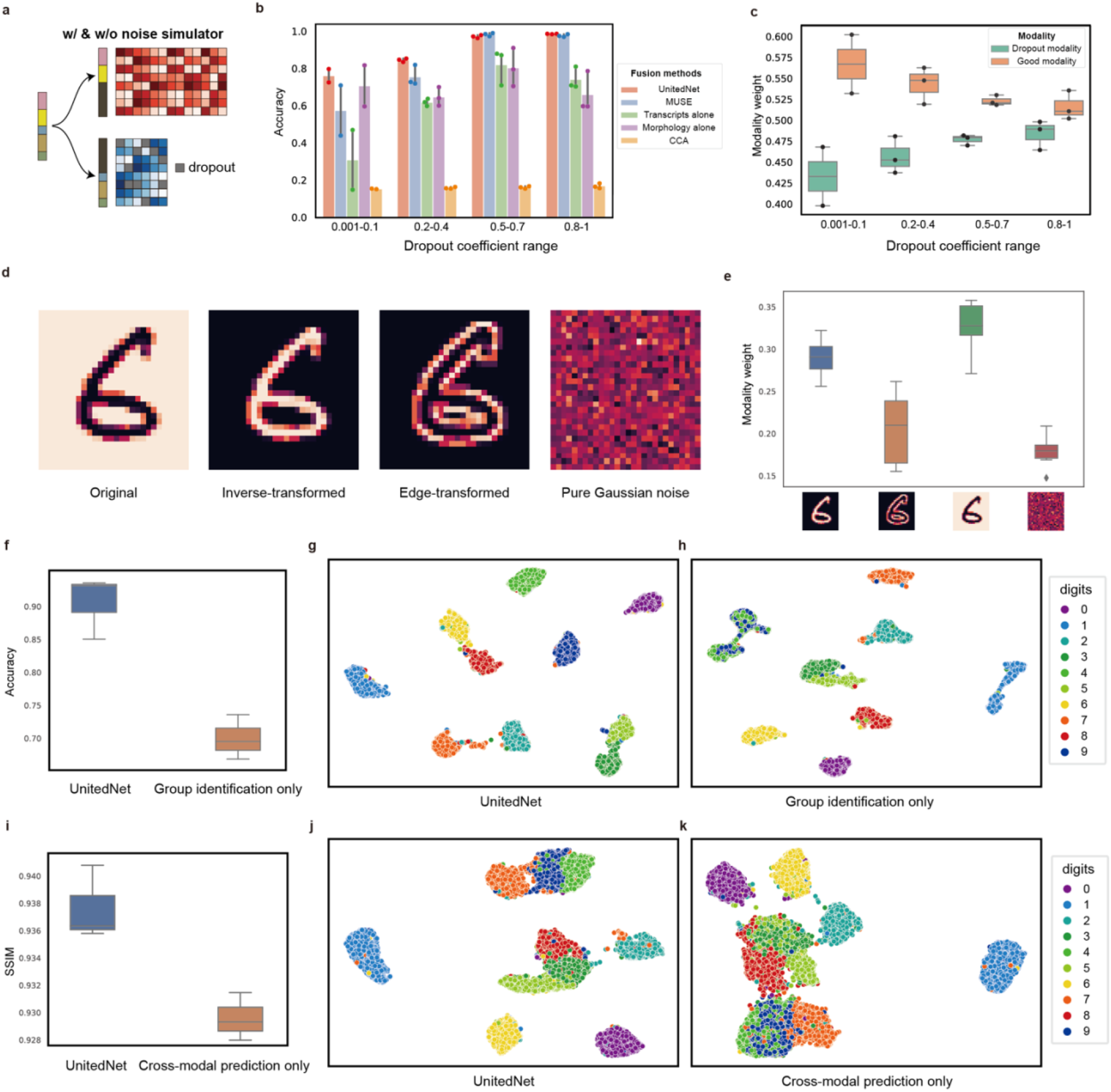
UnitedNet validation with a noised simulated dataset and a multi-modality MNIST dataset. **a**, Schematics of the MUSE simulator simulating the dropout effect in the transcriptomics data^24^. The transcriptomics modality is simulated with a controllable dropout level. **b**, The bar plot compares the accuracy of cell typing between UnitedNet and other methods as the dropout level changes. **c**, The box plot shows the modality weight learned in UnitedNet for different modalities of the simulated data in (a). **d**, An example of the four-modality MNIST handwritten dataset. We use the original, inverse-transformed, edged-transformed, and pure Gaussian noise as the four input modalities of UnitedNet. **e**, The bar plot shows the modality weight learned in UnitedNet for different modalities of the four-modality MNIST data in (d). **f**, The box plot compares the accuracy of unsupervised clustering for the MNIST data in (d) between the UnitedNet trained by both tasks and joint group identification task only. **g-h**, UMAP shows the shared latent codes of the MNIST data in (d) from UnitedNet trained by both tasks (g) and by joint group identification task only (h). **i**, The box plot compares the cross-modal prediction performance comparison between the UnitedNet trained by both tasks and the cross-modal prediction task only in terms of structural similarity index (SSIM) on the MNIST data in (d). **j-k**, UMAP shows the shared latent codes of data in (d) UnitedNet trained by both tasks (j) and by cross-modal prediction task only (k).

**Extended Data Fig. 4.**
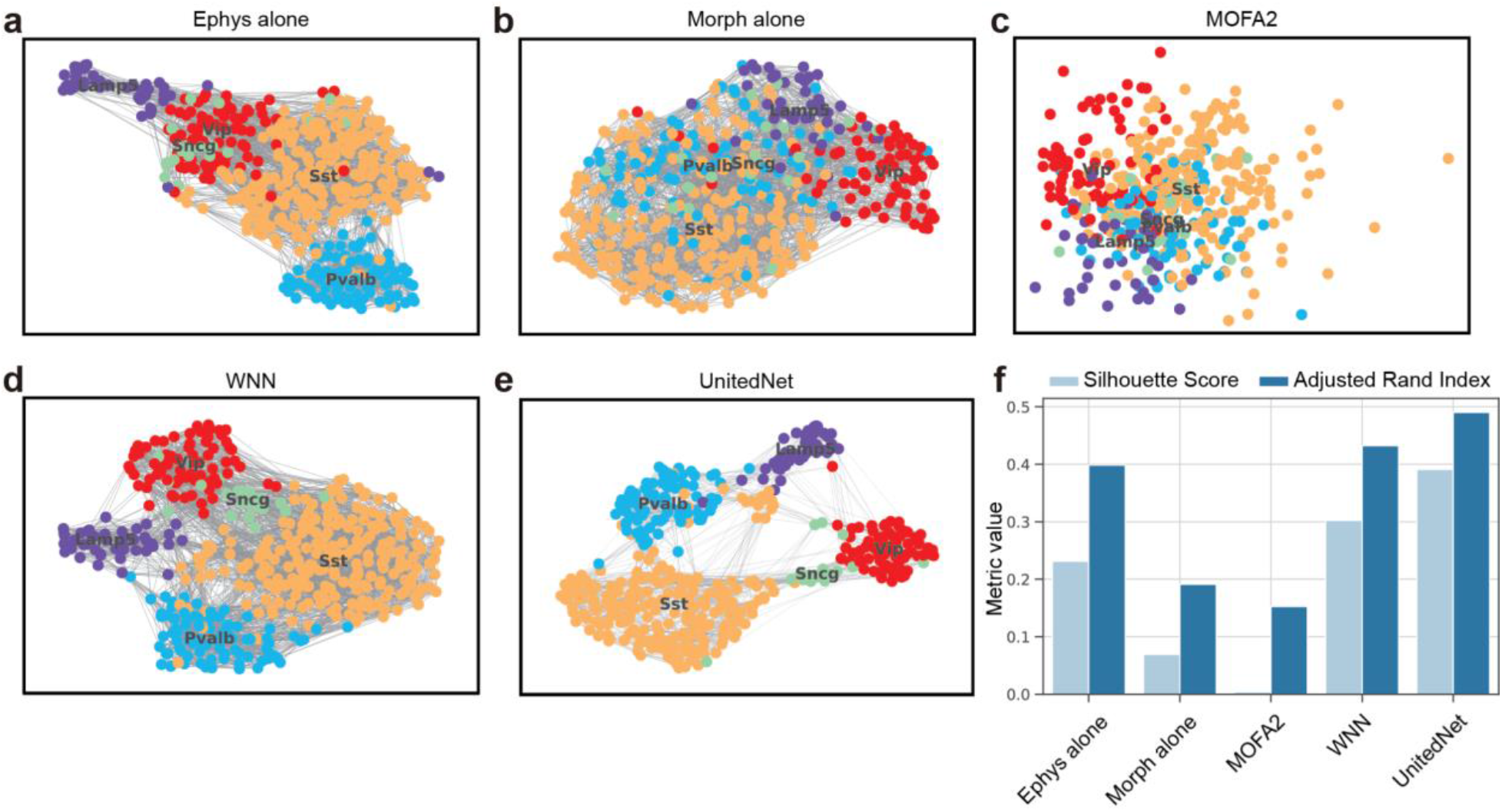
Fusion methods benchmarking on electrophysiological and morphological modalities from the Patch-seq dataset. **a-e**, The latent space visualization by UMAP of electrophysiology data alone (a), morphology alone (b), and shared/integrated codes from MOFA2 (c), WNN (d), and UnitedNet (e). The latent codes are colored to represent different cell type labels. **f**, The performance comparison of UnitedNet and methods in a-d on multi-modal fusion and unsupervised joint group identification. UnitedNet achieves the highest separability and identification accuracy in terms of the Silhouette score and adjusted rand index.

**Extended Data Fig. 5.**
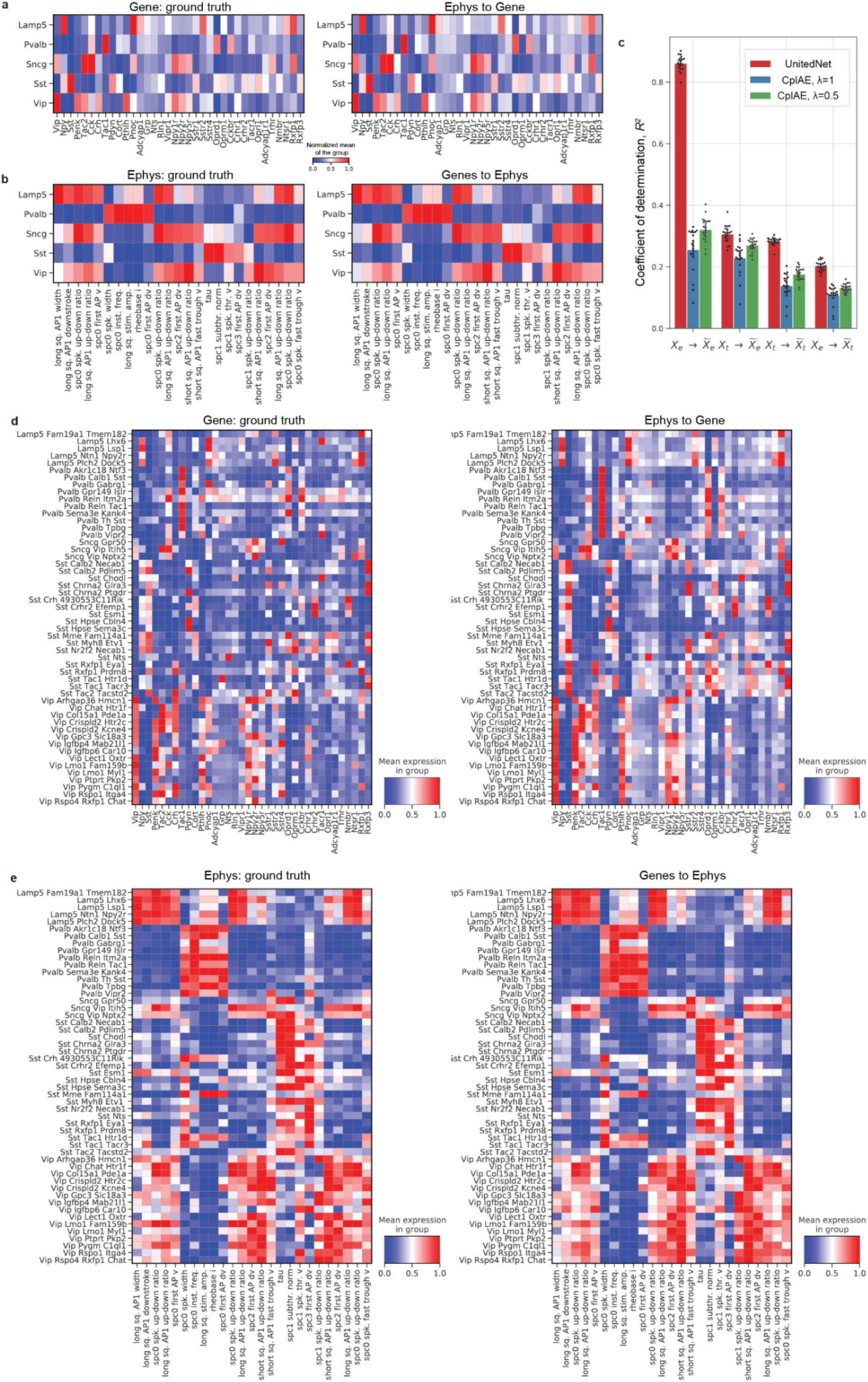
Cross-modal prediction result on electrophysiology and transcriptomics modality from the Patch-seq dataset. **a-b**, Heatmap comparing cross-modal predicted gene expression and electrophysiological features averaged over annotated major cell types with the ground truth. **c,** Bar and dot plots comparing within- and cross-modal prediction performances between two modalities (transcriptomics and electrophysiology) by coupled autoencoder (CplAE) with different values of hyperparameter λ and UnitedNet on 3,316 cells. Error bars show mean ± s.d. over 20-fold cross-validation. **d-e**, Heatmap comparing cross-modal predicted gene expression and electrophysiological features averaged over annotated detailed cell types with the ground truth.

**Extended Data Fig. 6.**
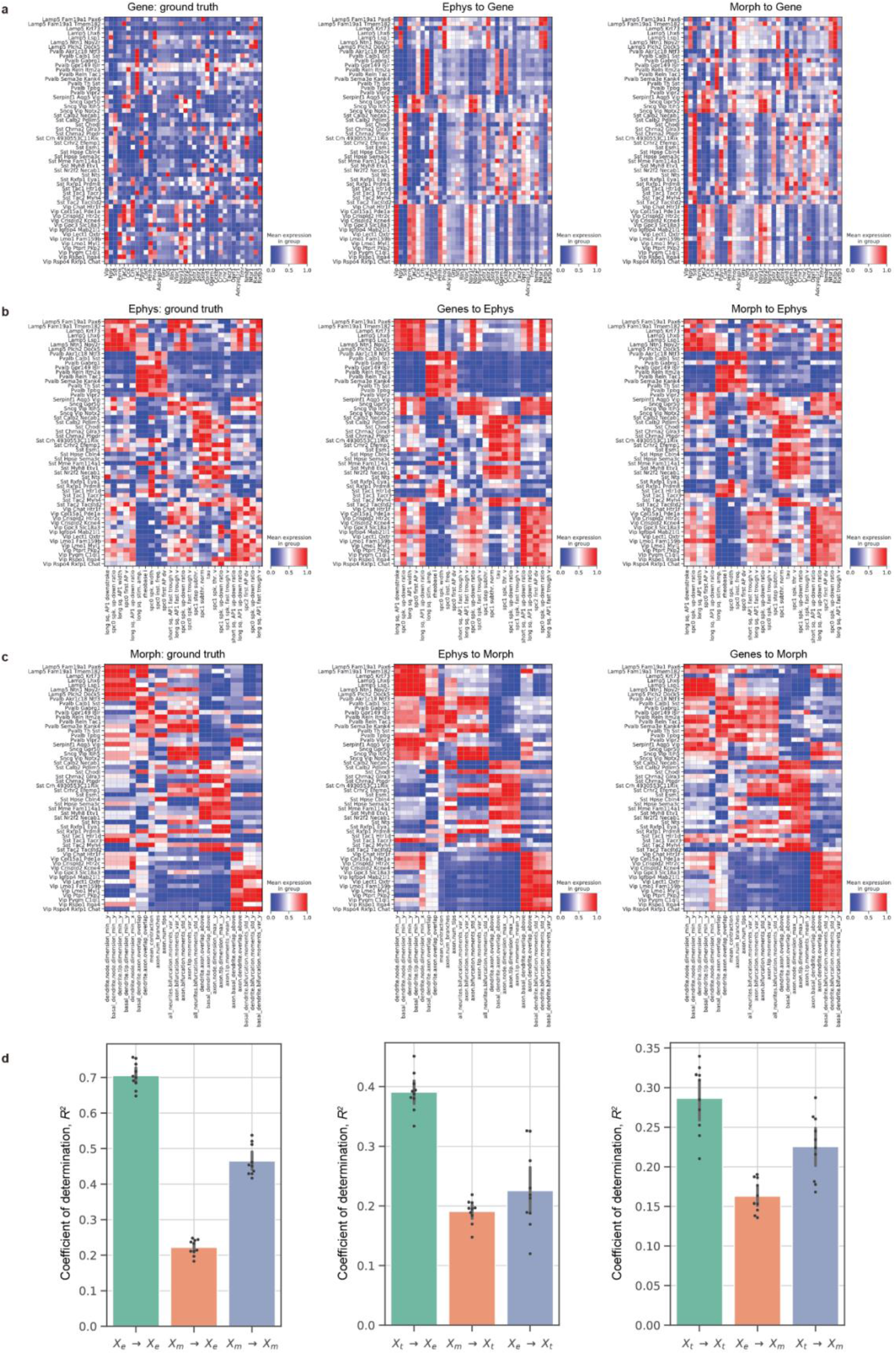
Translation results of Patch-seq with detailed cell type trained with E-T-M data. **a-c**, Heatmap comparing cross-modal predicted gene expression, electrophysiological and morphological features averaged over annotated major cell types with the ground truth. **d**, Bar and dot plots of within- and cross-modal prediction performances among three modalities (transcriptomics, morphology, and electrophysiology) by UnitedNet on 448 cells. Error bars show mean ± s.d. over 10-fold cross-validation.

**Extended Data Fig. 7.**
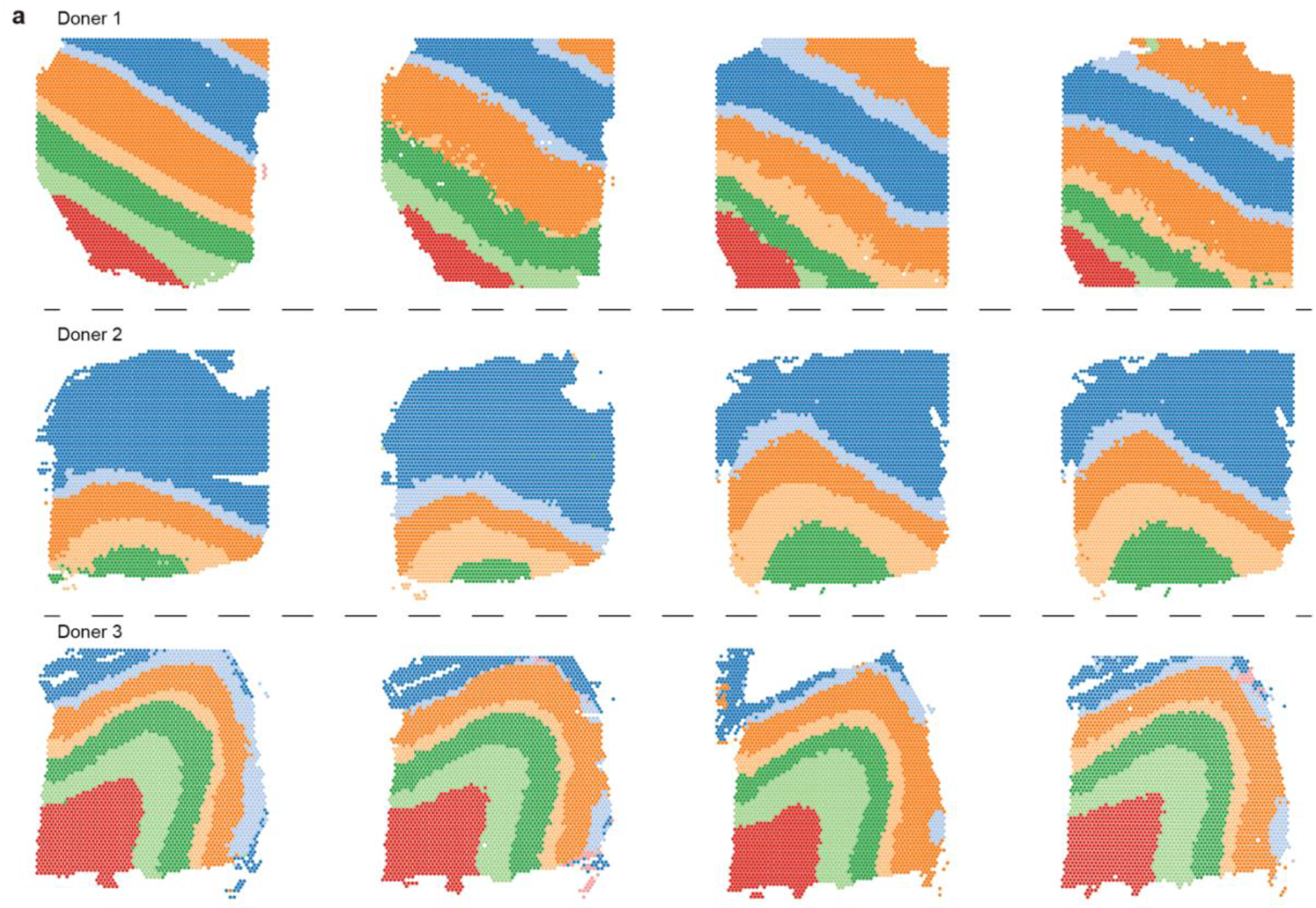
Annotation transferred result for DLPFC using UnitedNet. The spots of each measured brain slice are colored by tissue region labels predicted by UnitedNet. For each sample, UnitedNet is trained on the remaining 11 batches and is tested on this batch. Each UnitedNet successfully transfers the annotation of the reference brain slice region.

**Extended Data Fig. 8.**
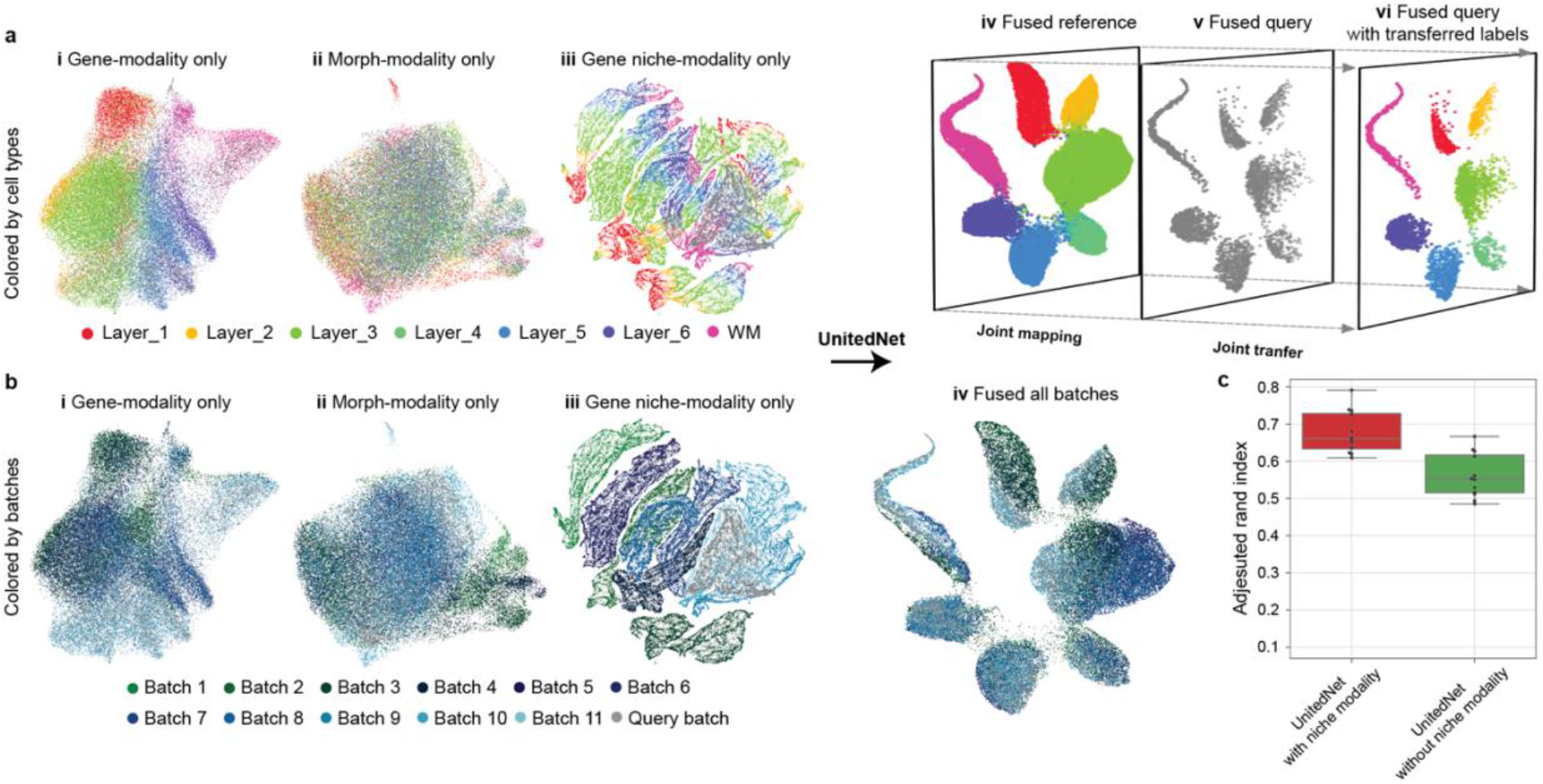
UnitedNet enables supervised group identification in the spatial-omics dataset. **a**, Supervised joint group identification enabled by UnitedNet. The UMAP latent space visualizations show gene expression (i), morphology (ii), gene expression niche (iii), and shared latent codes of UnitedNet (iv-vi) of the 12-batch DLPFC dataset. UnitedNet fuses the shared latent codes from three modalities and groups the codes based on cell type annotations (iv). Then, it projects the unlabeled query batch to the learned shared latent space (v), transferring the label of the reference to the shared latent codes of the unlabeled query batch to complete the annotation transfer (vi). The latent codes are colored by cell-type annotations. **b**, Joint batch effect removal enabled by UnitedNet. The UMAP latent space visualizations show gene expression (i), morphology (ii), gene expression niche (iii), and shared latent codes of UnitedNet of the 12-batch DLPFC dataset. The latent codes are colored by different batches. **c**, The boxplot shows the ablation analysis of cell niche modality. UnitedNet trained with gene expression, morphology, and cell niche modality shows better performance in joint group identification task than the UnitedNet trained with gene expression and morphology modality.

